# Engineering of interconnected microcapillary networks at the mesoscale via magnetic assembly of endothelial-cell ‘seeds’

**DOI:** 10.1101/2025.05.20.655072

**Authors:** Katarzyna O. Rojek, Antoni Wrzos, Fabio Maiullari, Konrad Giżyński, Maria Grazia Ceraolo, Claudia Bearzi, Roberto Rizzi, Piotr Szymczak, Jan Guzowski

## Abstract

Despite significant developments in endothelial-cell (EC) manipulation techniques, a proper *in vitro* model of a functional microvasculature with controlled local interconnectivity under well-defined global architecture is still lacking. Here, we report the generation of such controlled multi-scale vascular networks via manipulation of tens of sprouting EC ‘seeds’. We exploit magnetic patterning to assemble EC-coated superparamagnetic microbeads into ordered arrays and establish effective growth rules governing the development of interconnectivity and directionality of the networks depending on the applied seed-seed spacing. The EC-seed-based approach offers a range of advantages over conventional EC-manipulation techniques including: (i) expedited sprouting, (ii) spatial control over interconnections, (iii) reduction in cell consumption by >100x, and (iv) native high-throughput format. We co-develop multiparametric morphometric analysis tool and demonstrate high-content assessment of drug-induced vascular remodeling in 3D tumor microenvironments. Overall, we propose a uniquely precise and standardized vascular-microtissue engineering tool with applications, e.g., in angiogenesis research, high-throughput drug testing including personalized therapies, and with possible extension to organ-on-chip and tissue-regenerative approaches.

## INTRODUCTION

Precise modeling and design of human tissues or organs *in vitro* for the use in preclinical research or tissue regeneration requires incorporation of a functional vascular system, a branched network of channels enabling the supply of nutrients to the cultured cells, thus preventing necrosis, otherwise caused by hypoxia and malnutrition [1]. In the case of minimal tissues engineered for use in drug testing, it is critical to provide a microvasculature of well-controlled and reproducible morphology to warrant consistent results. In particular, the differences in the size and geometry of the capillary network may affect the kinetics of penetration of drugs inside the cultured tissue and thus affect the estimated drug efficacy. Currently, the most commonly used methods of producing tissues with embedded microvasculature rely on the spontaneous self-organization of endothelial cells (ECs) into vessels inside an external hydrogel matrix [2]. However, owing to the initial random dispersion of ECs in the hydrogel, the resulting microcapillary networks are also random and difficult to control [3–8]. Generation of such random vasculatures is also inefficient in terms of cell consumption and often results in a large fraction of disconnected or otherwise non-functional vessels [4, 5, 8].

Here, we demonstrate how the spatial organization of the microvasculature can be controlled at both microscopic and mesoscopic level via directed-assembly of EC-microcarriers, which we also refer to as EC ‘seeds’, into ordered quasi-2D arrays, giving rise to an interconnected vasculature of well-defined global architecture. We exploit magnetic hedgehog-like templates—arrays of permanent neodymium micromagnets arranged underneath a cell culture chamber—which guide the assembly of superparamagnetic microcarriers into a pre-designed quasi-2D pattern. Under culture, the EC-seeds ‘sprout’ and interconnect to form a percolated network. Compared to the previous EC-assembly methods [3, 9–11] our approach is unique in that—rather than manipulating *individual ECs*—it relies on the use of *EC microcarriers*. We use a numerical workflow that we introduced previously [12] to further develop a dedicated custom image processing software allowing multi-parametric morphological and topological characterization of the system of multiple sprouting and interconnecting microvasculatures.

The currently available methods of EC patterning, such as extrusion bioprinting [13–18] typically suffer from low reproducibility and limited angiogenic potential of the fabricated constructs. EC-spheroid-based bioprinting [19] provides further improvements, but also faces unresolved issues such as nozzle clogging or spheroid deformation and breakup during printing. More recent advances include aspiration-assisted bioprinting [20–23] which eliminates clogging but also limits the throughput. The most recent reports on parallelization [23] are promising in this respect, however, the use of EC-spheroids, overall, faces the issue of reproducibility associated with natural polydispersity of the spheroids, where the coefficient of variation (CV) of the spheroid diameter is typically in the range CV ∼ 10-20 % [24, 25].

A number of alternative approaches have been proposed, of which perhaps the most promising is the use of magnetically labelled ECs or magnetically ‘doped’ EC-aggregates and their magnetic assembly into quasi-2D patterns of controlled architecture [10, 26, 27]. However, the previous efforts have thus far provided rather small EC-assemblies with little sprouting potential. In addition, they also required the introduction of magnetic nanoparticles inside the cells or into the spheroids [9, 26]. Such invasive procedures are, in general, disadvantageous due to the possibility of altering cell behavior [26]. The magnetic assembly of single ECs are also excessively time-consuming, where aggregation of only around 100 cells typically takes up to several hours [26, 28]. Finally, the assembly of functional (e.g., lumenized) vascular networks of controlled architecture has not yet been demonstrated with any type of magnetic assembly approaches.

The use of EC-coated microbeads as EC-microcarriers have been proposed recently as a possible alternative route towards vascularization [29–31], but no specific methods of EC-carrier assembly into larger patterns have been proposed. Here, for the first time, we propose the use of superparamagnetic microparticles coated with EC-monolayers as the EC-seeds amenable to directed self-assembly into predesigned patterns. Microcarrier-based magnetic patterning provides for several advantages over other EC-pattering approaches including (i) the use of native (non-magnetically-modified) ECs (ii) reduced consumption of ECs, (iii) high precision of patterning warranted by the use of micromagnets and by high monodispersity of the microcarriers (CV ∼ 1-2 % in our case, see Methods section) (iv) high angiogenic potential of the EC-microcarriers resulting in rapid formation of well-defined vasculature with controlled local features (e.g. interconnections), and (v) the possibility of precise tracking of individual sprouts and their interconnections in time and space. Also, we use transparent polycarbonate (PC) plates to host the permanent micromagnets which allows for direct monitoring of the magnetic EC-seed assembly process *in situ* and facilitates manual correction of defects. Our micro-milled PC templates provide some practical advantages over the previously exercised ferromagnetic iron templates, so-called pin-holder devices [9]. In comparison, our method is simple and does not include the use of any expensive equipment such as a bioprinter or a discharge device (for iron microfabrication).

We use our system to establish a critical seed-seed spacing below which the neighboring microvasculatures become interconnected, and above which they remain disconnected, even at late times of culture. In the latter case, they can be treated as practically independent which allows systematic monitoring of multiple EC-seeds developing in nearly identical conditions. We show that the microvascular arrays co-cultured, e.g., with cancer cells (HeLa in our case), can efficiently serve as a high-throughput platform for functional high-content screening of various anti-angiogenic compounds in full 3D microenvironment [32]. In this respect, our system allows the measurement of a range of morphological and network-topological parameters unavailable with more conventional angiogenesis or vasculogenesis assays [33–36]. In particular, our high-content analyses are capable of differentiating between the impact of different drugs, here Taxol and Sorafenib—which we use as exemplary vascular endothelial growth factor (VEGF) or/and VEGF-receptor inhibiting agents (the former additionally being a cytostatic)—on the microvascular morphologies at various stages of development, i.e., just after sprouting or later after maturation and interconnection. Hence, in the future, our platform could be used to provide detailed insights into the mechanisms of action of drugs in various types of vascularized microenvironments.

We validate our method using both human umbilical vein endothelial cells (HUVECs) as well as human induced pluripotent stem cells (hiPSC)-derived endothelial cells. Via tight control over the bead-bead spacing, our method allows for the assembly of (i) ordered arrays of mature microcapillary networks which can be used as a robust high-content screening platform for testing of anticancer therapeutics or different anti- and pro-angiogenic agents, as well as (ii) interconnected capillary beds spanning the area of up to 1 cm x 1 cm (or possibly larger) for future applications, e.g., in regenerative medicine. Importantly, our platform could not only expedite the development of personalized treatments, but also serve as a standardized model system for studying the basics of angiogenesis or various microphysiological processes of healthy and diseased states involving cell migration—such as, e.g., metastasis.

## RESULTS

### Magnetic assembly of the EC-carrier microarrays

To achieve rapid and functional vascularization of hydrogel sheets with precise control over the vascular topology we assemble mesoscopic magnetic EC microcarriers (> 200 µm in diameter) into ordered arrays using external magnetic templates with well-defined and strong magnetic-field “hotspots”. Our system comprises two parts: (i) a PC plate with a micro-milled array of holes accommodating cylinder-shaped strong neodymium micromagnets and (ii) a culture chamber fabricated in polydimethylsiloxane (PDMS) with a bottom wall made of a sufficiently thin cover glass (Figure 1A, Supp. Figure 1A). During the process of magnetic assembly of the EC-microcarriers inside the PDMS chamber, the magnetic template is placed underneath the chamber and both parts are tightly aligned (Figure 1B). The EC-coated paramagnetic microparticles, serving as the EC-microcarriers, are then suspended in fibrinogen solution and poured into the PDMS culture chamber. We observe fast (lasting few seconds) spontaneous migration of the EC-coated microbeads towards the hotspots generated above the micromagnets, resulting in the formation of an ordered microbead-array (Figure 1C). We estimate the efficiency of this spontaneous assembly process to be around 80% meaning that we observe on the average around 80% occupied and 20% non-occupied hotspots; the misplaced beads are then manually corrected using titanium fine-tip tweezers (Supp. Figure 1B). In the future, one could imagine further automation of the process, e.g., via its combination with microcarrier-printing. However, even without such automated pre-positioning, our method has an important advantage in that the microbeads remain very efficiently immobilized at their equilibrium positions. This feature, in turn, allows reproducible generation of multiple (nearly) identical microbead-arrays using a single template, which is of great relevance in tissue modeling for use, e.g., in drug testing. In particular, such precise templating allows to differentiate between the intrinsic (biological) complexity of the system and the variations in system geometry.

**Figure 1.**
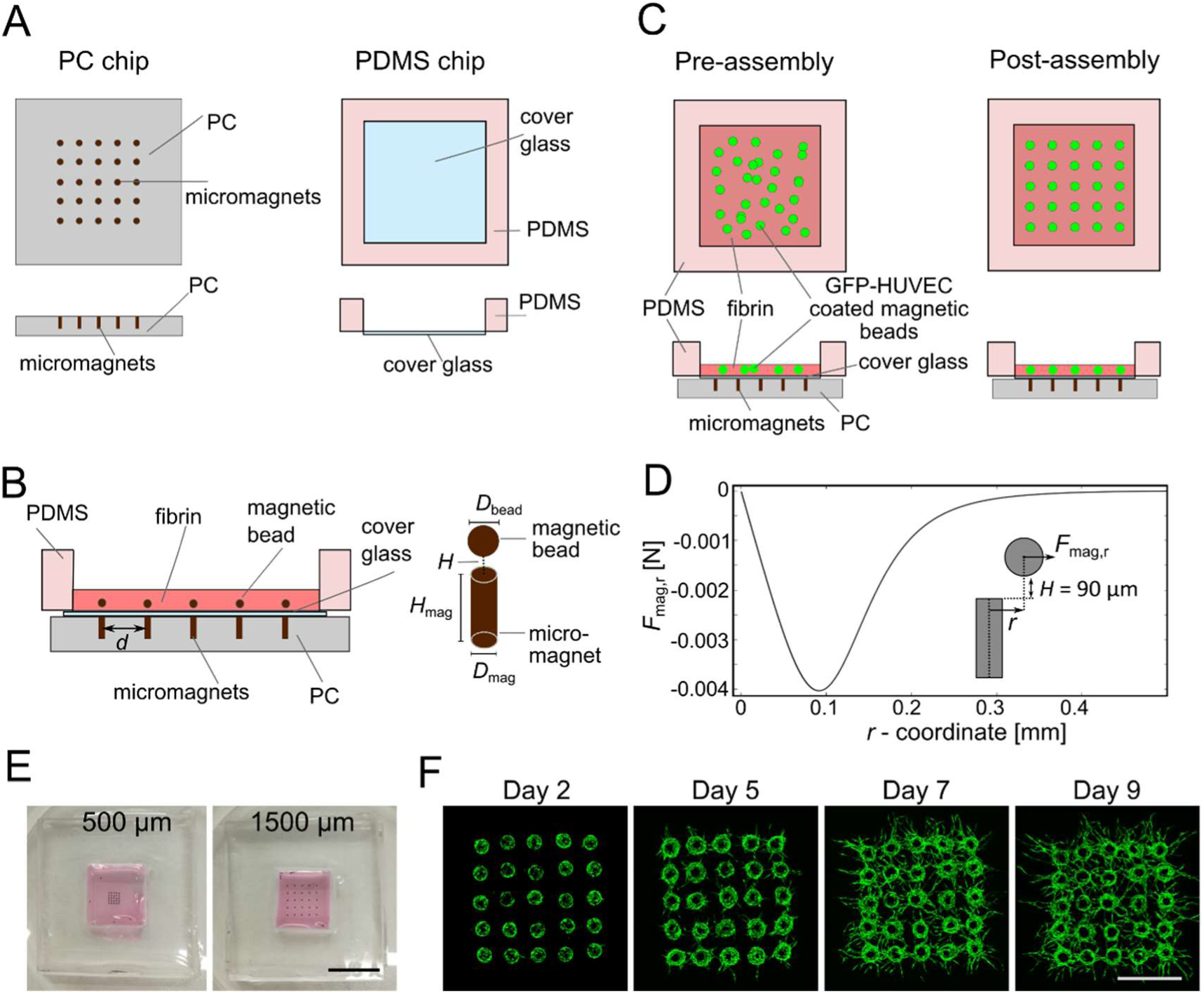
Generation of ordered arrays of microparticles over the substrate using magnetic-field patterning. **A)** Sketch of the experimental setup: a PDMS culture chamber with the glass bottom coverslip and a PC chip hosting the micromagnets. **B**) Illustration of the assembled device with the PDMS chip placed above the PC chip (left) and magnetic microbead localized precisely above the micromagnet. *d* – distance between the beads, *D*_bead_ – bead diameter, *H* – distance between the microbead and micromagnet, *H*_mag_ – micromagnet height, *D*_mag_ – micromagnet diameter. **C**) Scheme illustrating generation of ordered arrays of microparticles coated with endothelial cells. Left: PC chip hosting the micromagnets and the PDMS chamber filled with magnetic EC-coated microbeads suspended in non-crosslinked fibrin mixed with thrombin. Right: fibrin crosslinks after 10 mins leaving enough time for the magnetic self-assembly to take place. **D**) Results of the numerical calculation of the radial component F_mag,r_ of the magnetic force *F*_mag_ acting at a probe microparticle depending on its radial displacement *r* from the micromagnet axis for the experimentally relevant spacing *H* = 90 µm. Positive direction of the force is outward (i.e., negative values correspond to magnetic attraction). **E**) Images of PDSM culture chambers with EC-coated magnetic microbeads seeded at different densities in the fibrin hydrogel (spacing between the beads: 500 and 1500 µm respectively). Scale bar 1cm. **F**) Confocal images of the array of HUVEC-GFP-coated microbeads visualized at day 2, 5, 7 and 9 of culture. Scale bar 1000 µm.

We note that, in general, it is not obvious that the beads would necessarily tend to align perfectly with the micromagnets. In fact, the equilibrium XY-position of each bead may depend on the details of the shape of the micromagnet, as well as the distance in the Z-direction separating the bead and the micromagnet. To check the equilibrium positions of the beads in our case, we performed direct numerical calculations of the magnetic field distribution around a cylindrical micromagnet using COMSOL Multiphysics 6.1 with the AC/DC Module. We set the diameter of the microbead (*D*_bead_ = 265 µm) and the dimensions of the micromagnet (*D*_mag_ = 200 µm, *H*_mag_ = 500 µm) to reflect the experimental conditions. We also fixed the spacing *H* between the upper micromagnet surface and lower microparticle surface to be equal to the thickness of the cover glass used in the experiment, *H* = 90 µm. The results (Figure 1D) confirm that the radial component *F*_mag,r_ (in the XY-plane) of the magnetic force *F*_mag_ acting at a probe magnetic microparticle is always attractive, i.e., directed towards the symmetry axis of the micromagnet (the Z-axis), which accordingly sets a unique and well-defined equilibrium position of the microparticle to be at the Z-axis, i.e., at a radial distance *r* = 0 from the axis. We further also checked that this feature (attraction to the Z-axis) actually holds for all tested Z-spacings in the range *H* ∈ [50,110] µm corresponding to typical ultra-thin cover-glass thicknesses. Accordingly, the microparticle pattern assembled above multiple micromagnets can be expected to perfectly reproduce the micromagnet pattern. For comparison, we performed analogical calculations for the case with the magnetic hotspot generated by a ‘pin-holder’ device consisting of an iron micro-cylinder or ‘pin’, magnetized by another large permanent magnet placed underneath, as proposed previously by the Honda group [9]. In particular, in the case of small *H* values (below 70 µm), we found the equilibrium condition (zero radial force) to be not at the Z-axis but rather at a distance *r* = *r*_0_, with *r*_0_ depending on *H* (e.g*. r*_0_ ≍ 70 µm for *H* = 50 µm), that is along a circular region of radius *r*_0_ centered around the Z-axis. This means that in the case of magnetized pins the equilibrium positions may become ‘degenerated’, i.e., no longer unique but rather distributed anywhere along the circular regions above the pins (Supp. Figure 1C). In the case of a microparticle array assembled above such magnetized pins, this effect would lead to random perturbations in the spacing between the neighboring microparticles. Altogether, the use of permanent cylindrical micromagnets provides for highly precise patterning independent of *H*.

Next, we demonstrate experimentally that the system can be indeed used to precisely control the spacing between the EC-coated microparticles (Figure 1E). To this end, we fabricate several different magnetic templates with varying micromagnet spacing. We carefully move the assembled microparticle arrays into the cell culture incubator for 15 mins allowing the fibrin hydrogel to fully crosslink. Next, we remove the PC magnetic templates, place the PDMS culture chambers in the P60 petri dishes and culture the arrays for up to 2 weeks allowing EC-coated microbeads to produce vascular sprouts (Figure 1F). At pre-determined time-points (typically day 2, day 5, day 7, day 9), we remove the arrays from the incubator to acquire confocal fluorescence images of the arrays of HUVEC-coated microparticles. Noteworthy, imaging of the sprouts (which typically grow not at the bottom but deeper in the hydrogel) is facilitated via the use of an ultrathin (≤ 110 µm) bottom of the culture chamber. The hydrogel slabs hosting the arrays can also be easily isolated from the open-top PDMS chambers (Supp. Figure 1D) which demonstrates compatibility of our platform with downstream cell- and tissue characterization techniques (e.g., genomic analysis) which, however, we leave as future work, here focusing mostly on the morphometric and functional analyses.

### Initial distribution of EC-coated microbeads governs the final topology of microvascular networks

To demonstrate the application of microbead arrays in fabrication of spatially controlled and physiologically relevant microvasculature, we designed magnetic templates hosting the 5 x 5 arrays of micromagnets with a predetermined different micromagnet spacing *d*, i.e., *d* = 400, 500, 750, 1000 and 1500 µm, and studied the angiogenic sprouting behavior of the generated microvascular arrays (Figure 2A). We used GFP-expressing HUVECs (GFP HUVECs) as the vascular precursor cells, and normal human dermal fibroblasts (NHDFs) as the stromal cells, where the latter supported HUVEC morphogenesis, e.g., via secretion of pro-angiogenic growth factors and extracellular matrix proteins [35, 37]. To perform complex morphometric analysis of the sprouting networks we adapted and modified a custom image processing tool that we developed previously for the analysis of single sprouting beads [12]. The current modified software can not only automatically detect and analyze the sprouting behavior of the individual beads forming the *N* x *N* array but also detect the interconnections and analyze the directionality of sprouting (see Methods). Our image analysis workflow starts with the generation of a maximum intensity projection of an acquired confocal 3D scan, followed by segmentation of the image and its skeletonization necessary to compute the basic morphological and network-topological metrics such as: area, length and width of the sprouts, number of tips and primary branches, as well as the interconnectivity (Figure 2B). In particular, as the measure of the interconnectivity, we use (i) the total number of interconnections between the neighboring beads (Figure 2B, 2C.), and (ii) the total number of connected components, i.e., the number of interconnected groups of beads. We approximate the number of interconnections as the number of intersections of the sprouts with the borders of the Voronoi cell drawn around a given microcarrier under an additional requirement that the intersections are not along the tip segments of the sprouts. In such a way we eliminate the majority of sprouts that terminate just beyond the Voronoi border without reaching the neighboring bead (or anastomosing with its sprouts), i.e., those not forming the actual interconnections.

**Figure 2.**
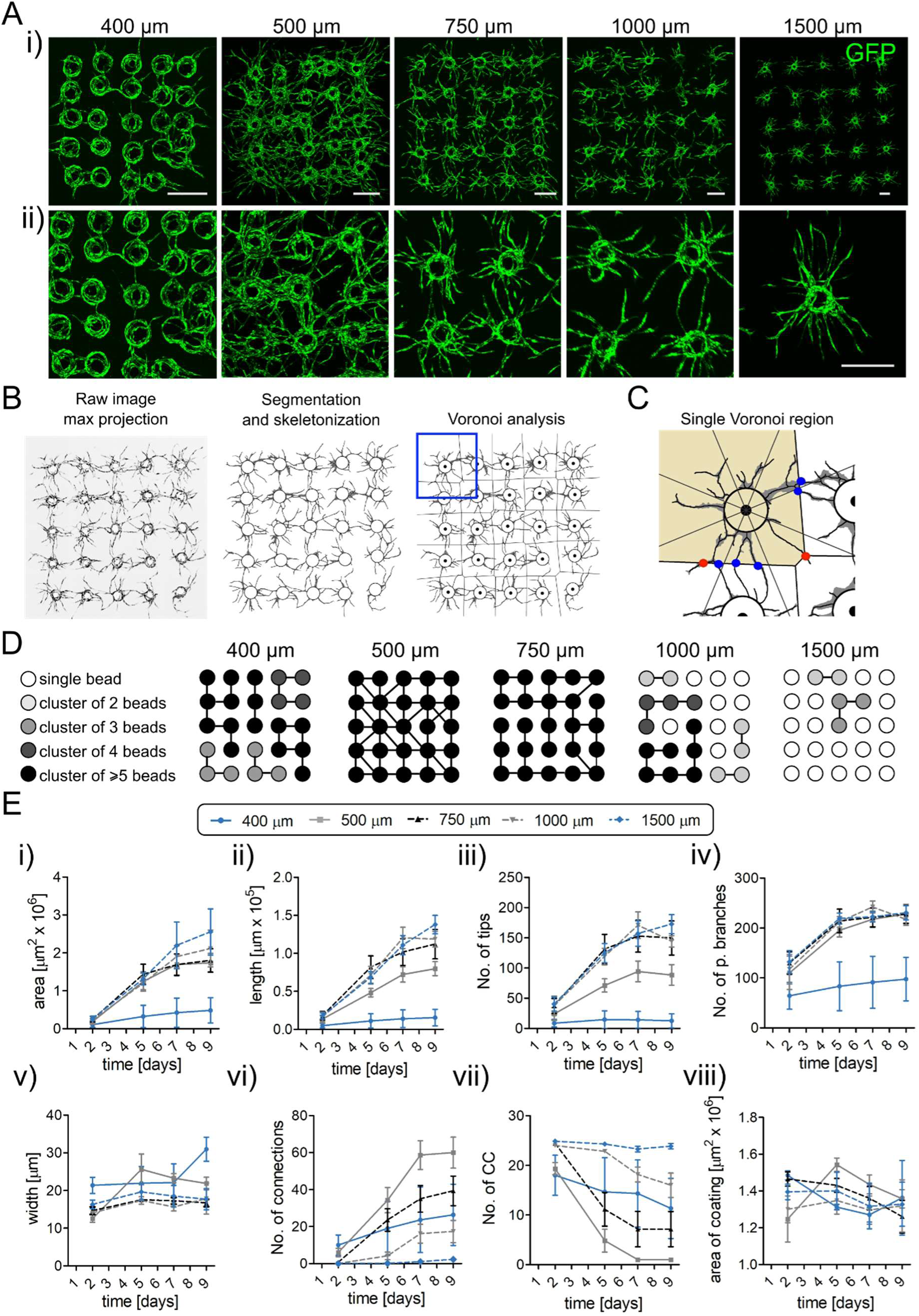
Distribution of the EC-coated microbeads in hydrogel (bead-bead spacing) affects the architecture and morphological properties of the microvascular networks. **A**) (i) Representative images of the arrays of microparticles coated with GFP-tagged HUVEC cells and seeded at different densities (*d* = 400, 500, 750, 1000 and 1500 µm respectively) imaged at day 7. Each array consists of 25 EC-coated microbeads of the same size (265 µm in diameter). (ii) High magnification images of arrays showing the differences in morphology of the developed microvascular networks depending on the spacing *d* between the microbeads. Scale bar 500 µm. **B**) Image processing workflow. From left to right: generation of the maximum projection image, segmentation and skeletonization, Voronoi analysis of connectivity. **C**) Illustration of a single Voronoi region. Points of interconnections between the beads are marked in blue. Points that intersect the Voronoi cell but do not result in a formation of interconnections are displayed in red. The image is divided into 8 zones to analyze the directionality of the vascular sprouting. **D)** Visualization of the number and size of bead-clusters connected via capillaries at various *d* as acquired from a single representative image for each *d* at day 7. Lines between the beads indicate the direct vascular connections. **E**) Morphometric analysis of the arrays: (i) total area of the sprouts, (ii) total length, (iii) number of tips, (iv) number of primary branches, (v) width of the capillaries, (vi) number of established vascular connections, (vii) number of connected components and (viii) area under the masks (i.e., the cells coating the beads). The numbers of the arrays in the assay (biological repetitions) were as follows: Spacing 400 µm, *n* = 3; 500 µm, *n* = 5; 750 µm, *n* = 7; 1000 µm, *n* = 6; 1500 µm, *n* = 7). Symbols in the graphs indicate the mean values, and error bars are the standard error of the mean (SEM). Statistical significance in **E** was analyzed using a two-way ANOVA with Bonferroni post-hoc test. See Supp. Table1 for detailed statistics for (**E**). Numerical values that underlie the graphs are shown in Supp. Data file.

We observed that the applied bead-to-bead spacing *d* in the array had a significant impact not only on the overall complexity of the network but also on the actual type of morphology of the developing endothelium (Figure 2A, 2D, 2E.). At the largest spacing, *d* = 1500 µm, *d*/*D*_bead_ ⋍ 5.7, we observed sprouting microvasculatures developing practically independently (Figure 2D), i.e., hardly showing any interconnections between the neighboring EC-seeds at the endpoint of the applied culture period (day 9). The measured dynamics of sprouting remained well aligned with the previously studied case of completely isolated beads, i.e., single beads cultured in separate wells [12]. At a smaller spacing of *d* = 1000 µm, yet still significantly exceeding the bead size, *d*/*D*_bead_ ⋍ 3.8, we observed occasional development of individual interconnections (Figure 2D). The total length and area of the sprouts in this case seemed slightly reduced as compared to the case *d* = 1500 µm, yet only at the later stages of development (days 7-9) (Figure 2E). At an intermediate spacing *d* = 750 µm, *d*/*D*_bead_ ⋍ 2.8, we observed a rapid increase in the number of interconnections, starting already at day 5, with an average of around 40 interconnections per matrix achieved at day 9, corresponding to exactly 1.0 interconnection per each pair of the neighboring beads (Figure 2E). We note, however, that not all of the nearest neighbors were interconnected in this case (Figure 2D). This is because some of the beads developed multiple interconnections with their neighbors while others remained disconnected. We observed further decrease in the area of the sprouts at days 7-9 (Figure 2E). At the spacing *d* = 500 µm, *d*/*D_bead_* ⋍ 1.9, we observed a maximum of 60 interconnections per matrix at day 9, that is around 1.5 interconnection per pair of neighboring beads (Figure 2E). In this case, we typically observed all the nearest neighbors to be interconnected, thus resulting in a fully interconnected vascular mesh (Figure 2D). At the same time, as compared to the cases with the larger spacings, we observed a significant decrease in the total length of the sprouts and an even more significant decrease in the number of free tips (Figure 2E). Since the number of primary branches remained unaltered for the bead spacings in the range *d* = 500 - 1500 µm (Figure 2E), the above result quantifies an important effect, namely that the developing of interconnections prevents further growth or branching around the interconnected regions. In fact, each interconnection corresponds to the elimination of a tip, while the growth (and branching) of the sprouts happens predominantly at the tips. Finally, at the smallest studied distances *d* = 400 µm, *d*/*D_bead_* ⋍ 1.5, we observed a significant change in the overall morphology of the developing endothelium (Figure 2E), which included merging of the endothelial cell monolayers (or multilayers) covering the neighboring beads, with scarce actual sprouts. The significantly reduced sprouting manifested itself, e.g., in a strongly diminished number of primary branches and free tips, even at the early stages of development (Figure 2E). Also, the total area of the sprouts and their total length were multi-fold reduced as compared to the cases with larger spacings (Figure 2E). In particular, the reduction factor for the area and length at day 9 were around 5 and 9, respectively, as compared to the case with *d* = 1500 µm (Figure 2E). The changes in the microvascular morphology in the case *d* = 400 µm can be attributed to the lack of available space for the development of the sprouts. This trend remains in line with the previously observed cases of EC cells growing in porous media, e.g., in the interstitial spaces between close-packed hydrogel microbeads [38].

Overall, upon decreasing the bead-bead spacing from *d* = 1500 µm down to *d* = 500 µm we find little change in the microvascular morphology, especially at early times of culture, up to day 5. At *d* = 500 µm, at later times, we observed more and more interconnections, reduced number of growing tips, and finally the formation of a fully percolated network. In fact, the case *d* = 500 µm appears critical in that it corresponds to the maximal observed connectivity, wherein further decrease in the spacing down to *d* = 400 um leads to severely suppressed sprouting and a tendency towards the formation of a space-filling endothelium.

Next, we re-analyzed the acquired confocal images in terms of the impact of the neighboring beads on the direction of angiogenic sprouting. We divided the beads into three subgroups depending on their position in the array: (i) ‘center’ (3 x 3 inner subarray, with each bead having 8 nearest neighbors), (ii) ‘edges’ (the beads at the sides of the array, corners excluded, each bead having 5 nearest neighbors) and (iii) ‘corners’ (each bead having 3 nearest neighbors), see Figure 3A, and we analyzed the dominant sprouting directions separately for each group depending on the bead-bead spacing *d*. In particular, we employed the Voronoi division and measured the mass distribution in the angular octants co-aligned with the principal- and diagonal directions of the (square) array, separately for each EC-coated microbead (Figure 3A). The results from the 4 different edge- and corner regions were pooled together, i.e., transformed to a common coordinate system. For edges, the principal direction (0 degrees) was set as perpendicular to the edge; for corners, as parallel to one of the neighboring edges and such that the matrix diagonal set the 45 degree direction, see Figure 3A. For the bead-bead spacings *d* = 400 µm and *d* = 500 µm we observed that, at day 7, in the edge and corner groups the octants with the largest endothelial mass were directed always *outside* the matrix (Figure 3A). For the microbeads located in the center of the array the directions with larger endothelial mass were the *diagonals* 45°, 135°, 225° and 315°, i.e., the directions pointing towards the further (diagonal) neighbors (Figure 3A). In contrast, for the spacings *d* = 750 µm and larger we observed isotropic distributions in all directions (Figure 3A). We additionally quantified this result via the coefficient of variation (CV), i.e., the standard deviation divided by the mean, of the probability values in different octants for various spacings and culture times (Figure 3B, Supp. Figure 2). As expected, for *d* ≥ 750 µm in all subgroups we found CV values around or below 10% at days 7-9 reflecting isotropic distributions. For *d* = 500 µm we observed strong anisotropy in the case of edge and corner beads, with CV around 35-40% at day 7 and 45-50% at day 9, but yet little anisotropy for the central beads (CV ∼ 15%). Similar or even slightly elevated values were observed for *d* = 400 µm. In addition, in the cases *d* = 400 µm and 500 µm we observed gradual growth of the anisotropy level at corners and edges during days 5-9 of culture. In contrast, for *d* ≥ 750 µm, we observed a relatively constant level at those times. In all studied cases, the elevated anisotropy levels at day 2 can be attributed to the small amount of sprouts at such early culture times.

**Figure 3.**
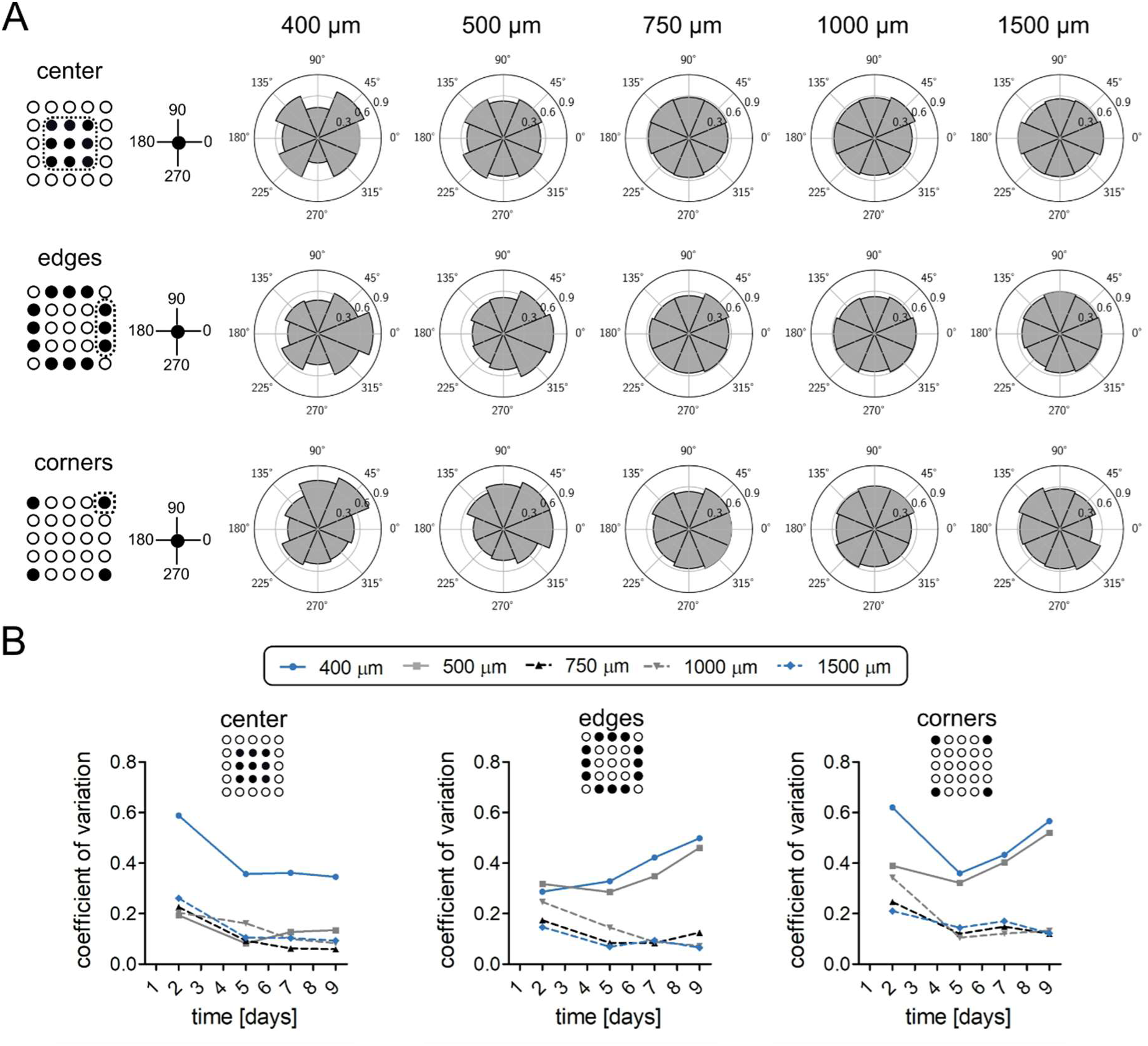
Analysis of directionality of the sprouting microcapillary networks. **A**) Radar charts illustrating the angular (azimuthal) distribution of the area occupied by capillaries at day 7 of culture. EC-coated microbeads were classified into 3 groups based on the number of neighbors’: center (8 neighbors), edges (5 neighbors) and corner (3 neighbors). Asymmetrical results from corner and bar regions were transformed to a common coordinate system, with all regions oriented to match the region indicated with a dotted frame (e.g. top bar is transformed by -90 degrees, etc.). The numbers of the arrays in the assay (biological repetitions) were as follows: Spacing 400 µm, *n* = 3; 500 µm, *n* = 5; 750 µm, *n* = 7; 1000 µm, *n* = 6; 1500 µm, *n* = 7). **B**) Coefficient of variation of the radar charts for EC-coated center-, edge- and corner microbeads at different indicated time-points, as a quantitative measure of the angular anisotropy of sprouting. Numerical values that underlie the graphs are shown in Supp. Data file.

Altogether, we find that the sprouts grow initially isotropically in all directions until the interconnections with the nearest neighbors (in the principal directions) are firmly established. In the case *d* = 500 µm, this starts to happen around day 5 (Figure 2E vi, 3B), after which the growth continues towards the regions of the remaining free space. Accordingly, the excess sprout area in those directions (Figure 3A) can be treated as an artifact of the anisotropy of the Voronoi cells. To eliminate this geometric bias, we additionally measured the angular dependence of *sprout coverage*. To this end, we calculated the area of the sprouts divided by the area of the Voronoi cell subsection in each octant. We evaluated this quantity for the central beads at day 7, corresponding to the region with an almost fully interconnected, mature network, where no (or little) further growth was observed. We found the sprout coverage to be again isotropic (Supp. Figure 3), which accordingly supports the picture of homogeneous penetration of all available space by the developing microvasculature until all interconnections have been established.

### Generation of complex geometries using magnetic force-based patterning of EC-coated microbeads

Vascularized tissue-specific models often require incorporation of microvasculature of a highly specific pre-defined architecture. To address this issue we microfabricated (PC)holders with several different micromagnet patterns (Figure 4A): (i) a hepatic lobule-like structure, (ii) a hierarchically branched vascular tree, (iii) parallel lines and (iv) a 3 x 3 array of empty squares (such structures could serve as arrays of ‘vascular beds’ [39]). Next, we used these magnetic templates to guide the assembly of HUVEC-GFP-coated microbeads. For the most efficient vascularization of the whole pattern, we used the optimal spacing *d* = 500 µm, warranting rapid development of interconnections between all (or nearly all) adjacent beads. Figure 4B shows the generated vascular structures imaged at day 1 and day 7 of culture. In the latter case, we observe systematic development of vascular interconnections between the neighboring beads which can be readily seen in the high-magnification images (Figure 4D). The demonstrated freedom of arranging the beads into arbitrary 2D patterns under well-defined and optimized adjacent bead-bead spacing constitutes an important step towards precision-engineering of interconnections in vascularized tissues, e.g., for future applications in perfusable organ-on-chip devices [5, 6, 40] or in fabrication of vascularized wound dressings.

**Figure 4.**
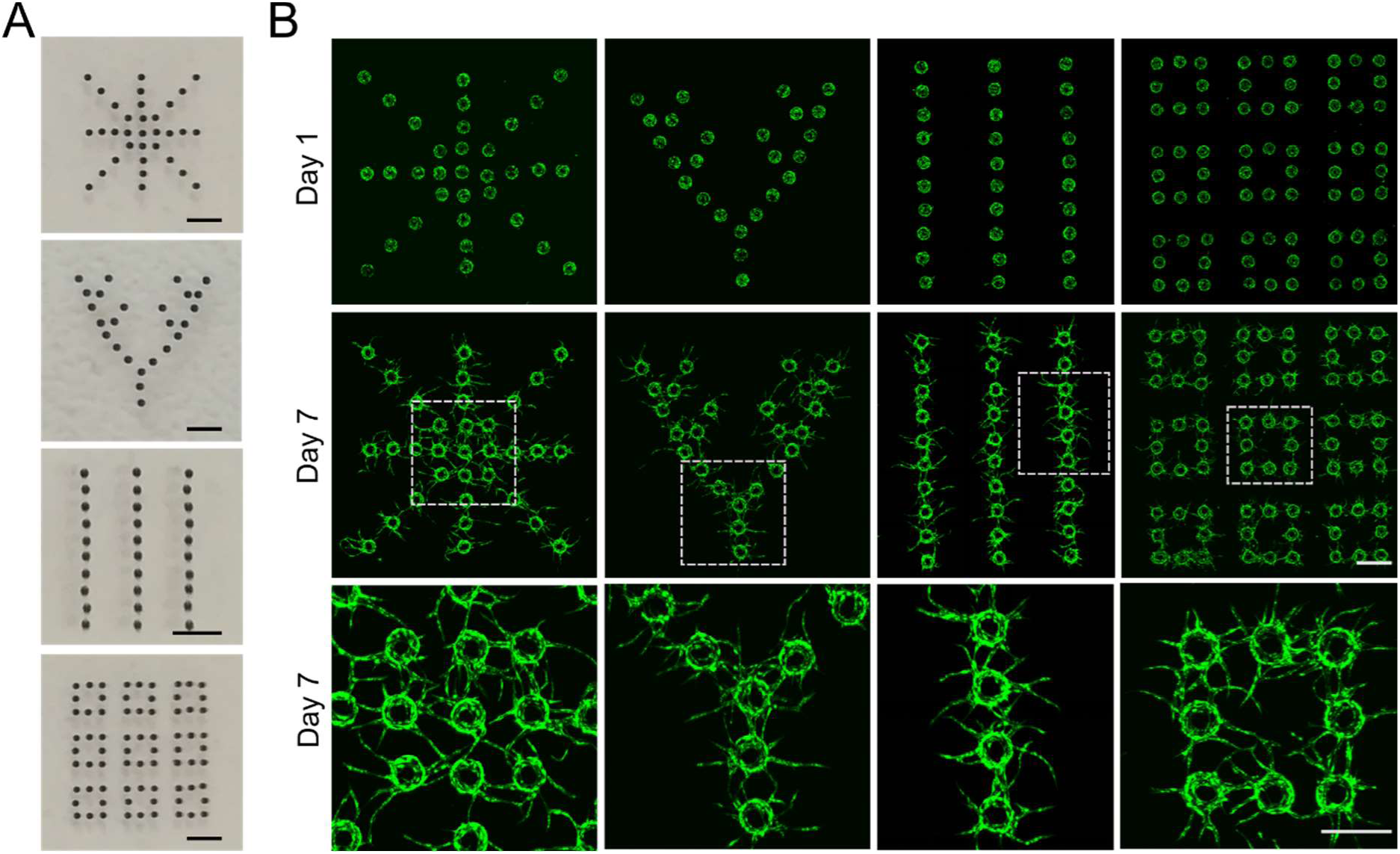
Complex patterns assembled from the EC-coated microbeads suspended in a fibrin hydrogel. **A**) Different magnetic templates used for the generation of different patterns. From the top: a model of liver lobule vasculature, a vascular tree with the hierarchical branching, parallel lines and an array of square-shape microcapillary beds. Scale bar 1000 um. **B**) Confocal images of GFP-HUVEC-coated microbeads patterned using different magnetic templates described above and imaged at day 1 and 7 of culture. Bottom panel shows high magnification images of the selected areas. Scale bar 500 um.

### Biological and morphological characterization of the engineered microvascular networks

To test whether the engineered microvascular networks exhibit proper physiological characteristics, we investigated the 3D structural integrity of the microvessels and verified the presence of characteristic marker proteins of healthy blood vessels. Immunofluorescence analysis showed abundant expression of CD31 (Figure 5A) and laminin (Figure 5B) in the microvascular arrays cultured for 7 days. The high magnification images (Figure 5A, 5B) visualizing the vascular connections between the HUVEC RFP-coated microbeads clearly display the mature morphology of the capillaries with the deposition of basement membrane by ECs around the perivascular extracellular matrix. Those results indicate vessel stabilization, high vessel integrity and characteristic elongated morphology of ECs co-aligned with the longitudinal direction of the capillary-like structures. Cross-sectional images of the engineered microcapillaries show the presence of a continuous lumen along the vessels, enclosed by ECs (Figure 5A, 5B). Moreover, we observed formation of apparently *toroidal* lumens around individual EC-coated microbeads (Figure 5C) and the formation of *tubular* vascular connections between those toroidal lumens (Figure 5D) resulting in the formation of a *continuously lumenized* structure. Last but not least, we found continuous cell–cell junctions lining the intersection of the ECs, as shown by the presence of the adherent junction protein, CD31 (Figure 5E). Overall, our results show that engineered arrays of microcapillary network-like structures supported by the microbeads display characteristic morphological and biochemical features of microvessels found *in vivo*.

**Figure 5.**
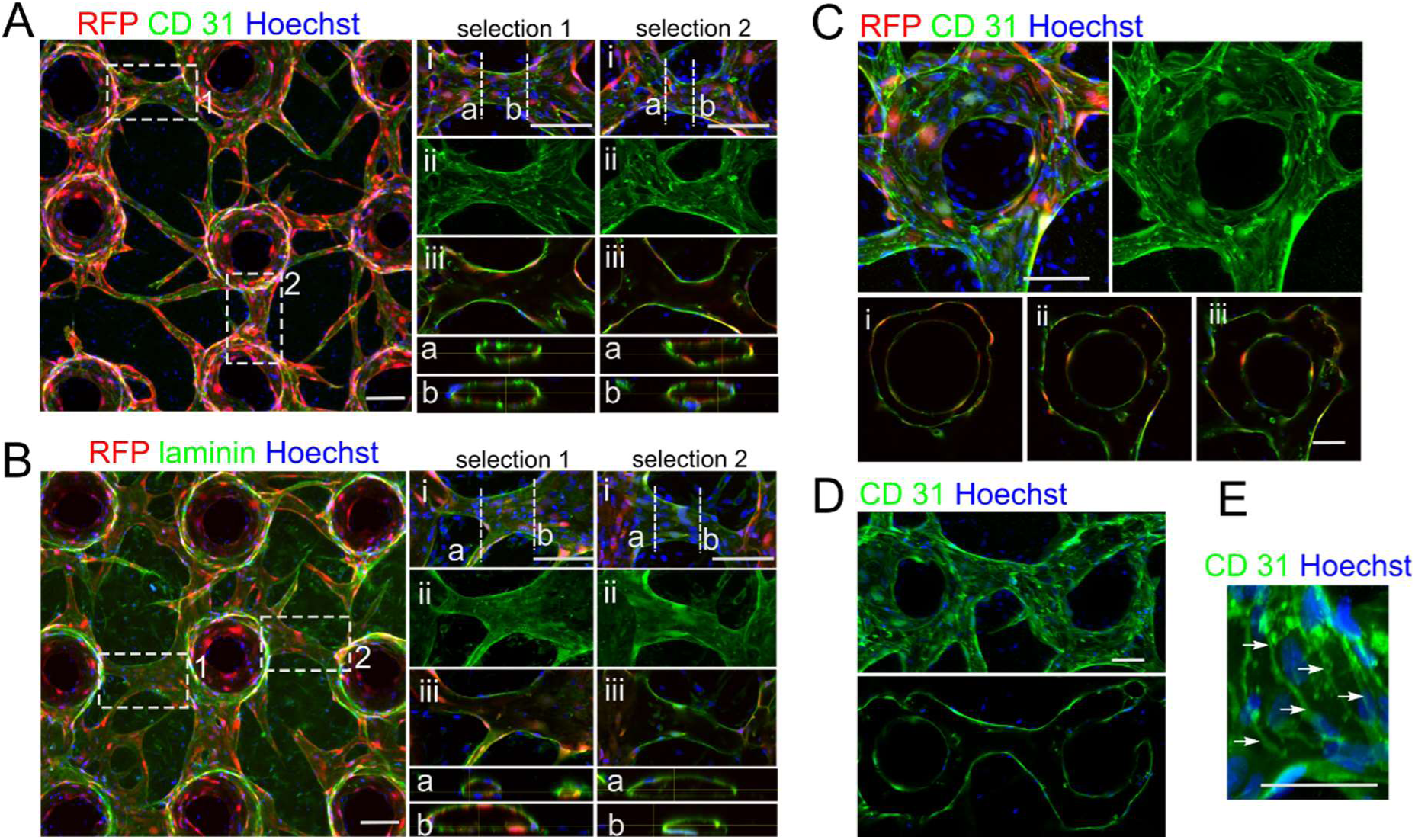
Arrays of HUVEC-coated microbeads form vessel-like structures characteristic of mature capillaries**. A-B)** Confocal images of arrays of HUVEC-RFP-coated microbeads seeded at 500 µm spacing and immunostained for (**A**) CD31 and (**B**) laminin. Right panels show high magnification images of selected areas (selection 1 and 2 i-iii) and their orthogonal cross-sections along the planes indicated with the dashed lines (a and b) showing lumen formation. **C**) Image of a representative HUVEC-RFP-coated microbead stained for CD31. Images in panels (i-iii) show the selected Z-slices visualizing the formation of lumens around the beads. **D**) Image of two neighboring HUVEC-coated microbeads immunostained for CD31. Bottom panel shows a single Z-slice illustrating the formation of a capillary tube connecting the two lumenized areas forming around the neighboring microbeads. **E**) Formation of tight junctions (indicated with arrows) between the HUVEC cells forming capillaries in the microbead-array assay. In all panels (**A-E**) Hoechst dye was used to visualize nuclei. Scale bar 100 µm.

### EC-coated beads accelerate the formation of microvascular networks

Next, we benchmarked the performance of the magnetically assembled arrays of EC-coated microcarriers (*d* = 500 µm) against the situation with ECs randomly dispersed in the fibrin gel, where the latter remains the most common method of producing microvasculature *in vitro* [1]. Our goal was to establish whether the strategy based on the use of microvascular ‘seeds’ can accelerate the formation of microvascular networks as compared to the more conventional approach, and what are the possible morphological differences in the generated capillary networks. First, we calculated the average number of HUVECs on the cell-coated beads (Figure 6A, 6B) and used the same number of ECs (per hydrogel volume) in both assays. Next, we imaged the growing vascular networks in both cases at day 2, 5 and 7 of culture. We observed that, after a week of culture, the arrays of EC-coated microbeads formed significantly larger and more complex vascular networks (Figure 6C). In particular, we found higher values of microvascular area and increased width of capillaries (Figure 6D). In the assay with dispersed ECs the width of capillaries did not change during the 7 days of culture, which may be attributed to slow cell aggregation and, in particular, to the lack of any pre-formed cell aggregates. As a result, the cells remained dispersed and did not form any bigger vascular structures. To quantify the complexity of the microvascular networks in the two cases, we defined the connectivity coefficient Con(*t*) as the decrease in the total number of connected components *N*_CC_(*t*) normalized to the situation with final fully connected network, that is we defined Con(*t*) = [*N*_CC_(0) - *N*_CC_(*t*)]/[ *N*_CC_(0) – 1]. In the case of EC-coated microbead arrays we found a rapid increase in Con(*t*), reaching the maximal values close to Con(*t*) = 1 already at day 5 of culture, corresponding to the formation of a fully interconnected vascular network (Figure 6D). In the case of dispersed ECs the values of Con(*t*) remained close to 0 at all times. Finally, to quantify the degree of cell aggregation, we also introduced the aggregation coefficient Agg(*t*), defined as Agg(*t*) = *A*(*t*)/*P*(*t*)^2^ where *A*(*t*) and *P*(*t*) refer to the area and the perimeter of the microvascular network, respectively. As expected, we found significantly higher values of Agg(*t*) for the microvasculatures formed by the EC-coated beads as compared to those formed by dispersed ECs, with the relative difference being up to 2 orders of magnitude at day 9 (Figure 6C).

**Figure 6.**
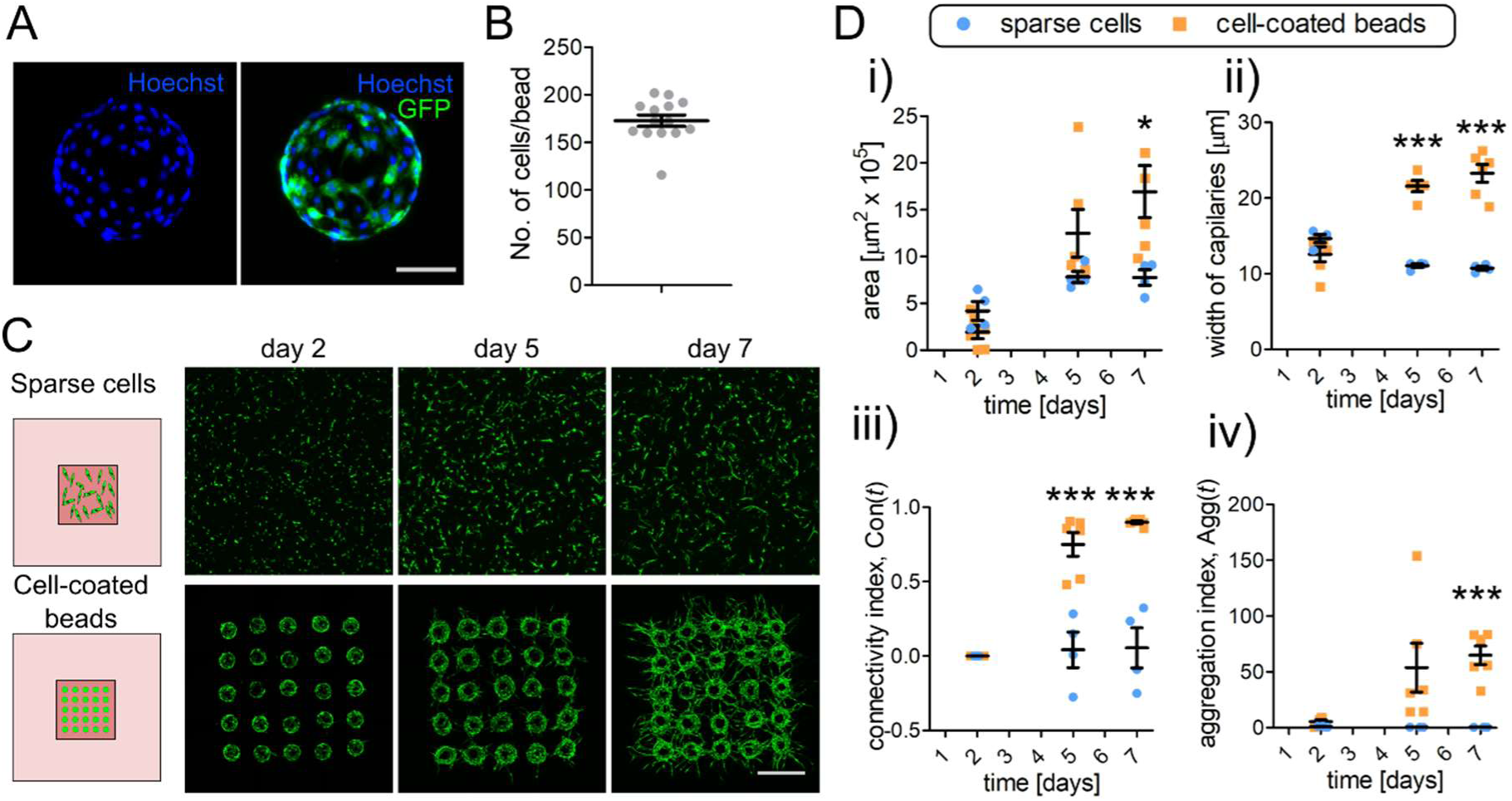
EC-coated microbeads accelerate the formation of vascular networks in 3D in vitro models. **A)** Representative image of a single GFP-HUVEC-coated microbead. Cell nuclei were visualized with Hoechst dye. Scale bar 100 µm. **B**) Quantification of the number of HUVECs used to coat one microbead (*n* = 14). Error bars indicate mean ± SEM. **C**) Images illustrating vascular network formation using conventional 3D in vitro assay where ECs are randomly dispersed in external hydrogel, here fibrin, and a microbead array model (bottom panel). In both cases the initial number of HUVECs (per hydrogel volume) was the same. Confocal images were acquired at days 2, 5 and 7 of culture. Scale bar 500 µm. **D**) Analysis of the (i) the total area , (ii) the width of the capillaries, (iii) the connectivity index, Con(*t*) and (iv) the aggregation index, Agg(*t*) of the arrays of HUVEC-coated beads seeded at 500 µm spacing and dispersed HUVECs as described in C). Dispersed cells, *n* = 4; cell-coated beads *n* = 6. * *p* < 0.05, *** *p* < 0.001. Statistical significance was analyzed using a two-tailed unpaired t-test. Numerical values that underlie the graphs are shown in Supp. Data file.

We believe that the slow development of the microvasculature in the case of dispersed ECs was a result of scarce cell-cell interactions caused by low initial cell concentration in the hydrogel and in the absence of any pre-assembled EC monolayers. According to the literature, this strategy is successful only at sufficiently high concentration of the ECs in the hydrogel, i.e., of the order 0.5–2×10^7^ cells/mL [5, 8, 41]. In our experiments, we used 0.35×10^5^ cells/mL, which reflects the number of cells coating the microbeads in 5 × 5 microbead arrays. This demonstrates that the approach based on the use of microcarriers as microvascular “seeds” allows for the formation of highly interconnected microvascular networks with endothelial cell consumption around 2 orders of magnitude lower as compared to the conventional approach based on the dispersed ECs.

### Drug screening studies

To demonstrate the potential use of our device in anti-cancer and anti-angiogenic drug screening studies, we co-seeded fluorescently tagged human cervical cancer cells HeLa GFP with EC-coated microbeads and stromal cells (fibroblasts) thus creating a cancer-like microenvironment. We used HUVEC RFP to enable co-visualization of both cancer cells and ECs in the system. The microvascular arrays were allowed to develop for 4-5 days prior to drug exposure and then were treated with different concentrations of either Taxol (100 or 1000 nM) or Sorafenib (1 or 10 µM). Both drugs are known to affect vasculature, however they exhibit different mechanisms of action [42, 43]. Taxol affects microtubule dynamics and inhibits EC proliferation and migration. It also has been shown to downregulate VEGF expression [43]. Sorafenib is a tyrosine kinase inhibitor and acts on signal transduction pathways such as Ras-Raf-MEK-ERK pathway [42]. As the control, we used arrays subject to the concentration of DMSO corresponding to the amount of DMSO used in the applied doses of the drugs.

We used microvascular arrays consisting of either (i) well-separated microvasculatures (*d* = 1500 µm) (Figure 7A) or (ii) those forming a fully interconnected network (*d* = 500 µm) (Figure 7C). Such two-way approach enables the assessment of the action of the drugs both in terms of (i) inhibition of the sprouting potential as well as (ii) disintegration of the existing vasculature. We analyzed the acquired images of the arrays before the drug treatment (time *t*_0_; that is day 4 or 5 of culture depending on experiment) and at 48 h post-treatment (time *t*_1_) (Figure 7C, 7D, Supp. Figure 4) and quantified the relative change in the macrovascular network area, length, number of tips, width of capillaries, number of connected components and number of primary branches, as well as (in the cases with *d* = 500 µm) network connectivity.

**Figure 7.**
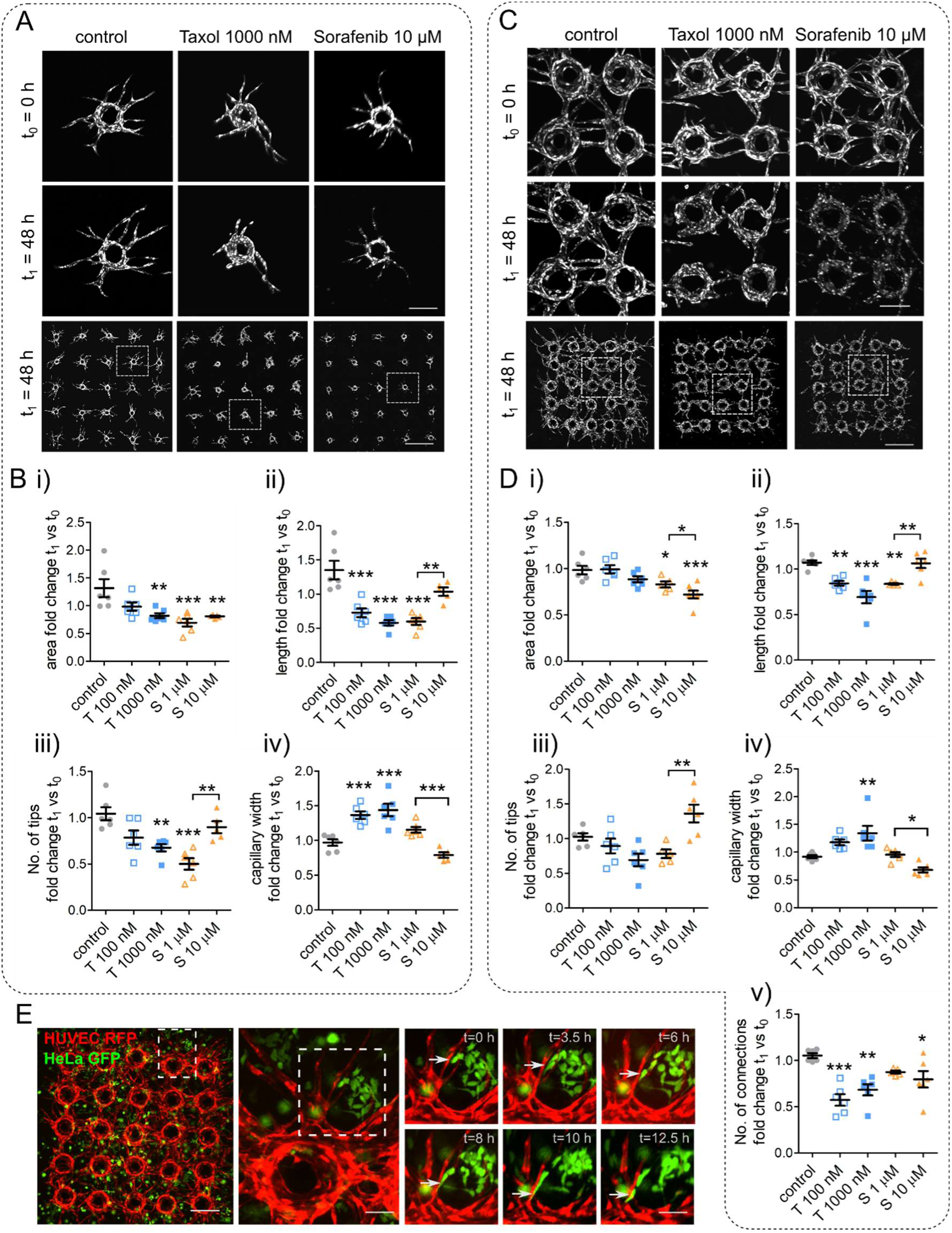
Application of microvascular arrays in anticancer drug screening studies. **A, C**) Images of the arrays of fluorescently tagged HUVEC-coated microbeads co-cultured with HeLa cells (not visualized) and seeded at (**A**) *d* = 1500 µm and (**C**) *d* = 500 µm spacing. At day 5 Taxol (100 or 1000 nM) or Sorafenib (1 or 10 µM) were administered to the structures; *t*_0_ – day 5 of culture, before drug exposure; *t*_1_ – day 7 of culture, 48 h after drug application. High magnification images (upper and middle panels) show selected areas of the arrays just before drug application (t_0_) and 48 h after drug exposure (t_1_). Scale bars 250 µm. Bottom panels show whole arrays. Scale bar 1500 µm in A and 500 µm in C. **B**) Morphometric analysis of the fold change in (i) the total area, (ii) the total length, (iii) the number of tips, and (iv) the width of capillaries at *t*_1_ relative to *t*_0_ for *d* = 1500 µm. Control, *n* = 6; Taxol (T) 100 nM, *n* = 6; Taxol (T) 1000 nM, *n* = 6; Sorafenib (S) 1 µM, *n* = 6; Sorafenib (S) 10 µM, *n* = 5. **D**) Morphometric analysis of the fold-change in (i) the total area, (ii) the total length, (iii) the number of tips, (iv) the width of capillaries and (v) the number of connections at *t*_1_ relative to *t*_0_ for *d* = 500 µm. Control, *n* = 6; Taxol (T) 100 nM, *n* = 6; Taxol (T) 1000 nM, *n* = 6; Sorafenib (S) 1 µM, *n* = 5; Sorafenib (S) 10 µM, *n* = 6. Error bars around the mean value indicate the standard error of the mean (SEM). * *p* < 0.05, ** *p* < 0.01, *** *p* < 0.001. **E**) Selected frames of the arrays of RFP-tagged HUVEC-coated microbeads co-cultured with HeLa GFP cells illustrating the migration of cancer cells into the microcapillaries. For full movies see Supplementary video 1 and 2. Scale bar 250 µm, high magnification images 50 µm. Statistical significance was analyzed using one-way ANOVA with Turkey post hoc test. Numerical values that underlie the graphs are shown in Supp. Data file.

In general, the two tested compounds affected the morphology of microvascular networks in a slightly different manner.

In the case *d* = 1500 µm, as shown in Figure 7C, we observed that the treatment with both drugs, applied at the highest concentrations, resulted in a significant reduction of the total area of the sprouts (Figure 7C) as compared to control. In the case of Taxol, we also observed a significant reduction of the length (Figure 7C, 7D) at both lower and higher concentration. However, in the case of Sorafenib we observed a rather peculiar effect in that the lower concentration (1 µM) had apparently much stronger effect on the length than the higher concentration (10 µM). Regarding the number of tips (Figures 7B, 7D), only the higher dose of Taxol and lower dose of Sorafenib led to the expected reduction. In general, the reduction in the number of tips can be interpreted as the effect of either tip retraction or less frequent branching (or both). However, again, the effect was absent in the case of higher dose of Sorafenib. The apparently weaker effects of Sorafenib at the higher dose can be explained in terms of a very rapid capillary disintegration, in fact happening so fast that the cells have no time to migrate, thus effectively ‘quenching’ the skeleton, as acquired from image processing. We stress however that the effect on the area in this case remains very strong (*p* < 0.01). Next, we also found the number of primary branches to be unaffected by the drugs, at least at the significance level *p* < 0.05, at any concentration (Supp. Figure 4C, 4D) This is reasonable as the drugs were administered only after the primary branches were already formed, i.e., when the number of primary branches normally reached a plateau (Figure 2E).

In the case *d* = 500 µm, as shown in Figure 7D, the results for the area and length were similar to the case *d* = 1500 µm (Figure 7B). In the case of Sorafenib treatments, we also observed higher numbers of tips as compared to the case *d* = 1500 µm (Figure 7B). We can attribute this effect to the drug-induced disintegration or ‘cutting’ of the pre-existing capillary connections. Further, similar to the case *d* = 1500 µm, we observed no significant effect on the number of primary branches (Supp. Figure 4D), except for the case of the higher dose of Sorafenib for which we found a significant increase as compared to control. Again, we interpret this result as an artifact of endothelial cell damage and the associated protein (GFP) aggregation, resulting in the numerical classification of the detected morphological irregularities as additional ‘sprouts’. Finally, as expected, we found that all treatments led to the reduction in the number of connections (Figure 7D) and a change in the number of connected components (Supp. Figure 4B), a signature of vessel ‘cutting’ and/or retraction. The effect was significantly stronger in the case of Taxol, for which we observed that the retraction of EC sprouts led to significant increase of the width of the sprouts and clearly visible retraction-bulb phenotype (Figure 7B, 7D).

In this study, we did not investigate the effect of Taxol and Sorafenib on cancer HeLa cell proliferation and viability in much detail. Nevertheless, we performed a basic analysis of the cancer cell mass approximated via the projected area of the visible cells. This parameter showed no profound effect on the cancer-cell number after 48h of treatment (Supp. Figure 4E, 4F), apart from a visible reduction in the case of Sorafenib (10 µM). Those results are in line with the previous results obtained in 3D models [6, 40, 44–46].

Finally, we also demonstrate that our model can be used to study the migration of cancer cells inside the microcapillaries in real-time. For this purpose, we performed time-lapse imaging of an array of HUVEC-RFP-coated microbeads co-seeded with HeLa GFP cells (Figure 7E). We observed that individual cancer cells often migrate into close proximity to the neighboring capillaries, trying to integrate with the microvascular network (Figure 7E). The possibility of direct tracking of cancer cells migrating inside (or outside) the capillary vessels in realistic microenvironments may be of significant interest to studies focusing on the metastatic potential of cancers forming solid tumors. The platform can be also applied to study the migration behavior of non-cancer cells, including tracking of individual endothelial cells or their different populations (Supp. Figure 5) to assess e.g. their mobility or/and sprouting phenotype.

In conclusion, our platform can be successfully used for the screening of drugs in well-controlled cancer microenvironments, providing (i) detailed and robust readouts in terms of the anti-angiogenic effects acquired from morphological changes in the tumor-associated microvasculature and (ii) insights into the interactions between tumor cells and the associated vascular network.

### Validation of the model with hiPSC

The recent advances in iPSC technology have made patient- and disease-specific human cells widely available as the sources for personalized drug screening tests, toxicology studies or disease models. To assess the future potential of our angiogenesis assay in personalized precision medicine, we also generated the arrays of EC-coated microparticles using hiPSC-derived ECs (hiPSC-ECs). Commercially available hiPSCs were first differentiated to an intermediate mesoderm stage and then to the final endothelial lineage (Figure 8A). We performed flow cytometry (Figure 8B) and immunofluorescence (Figure 8C) analysis on day 7 of the protocol to evaluate the EC differentiation. We observed that approximately 80% of cells were positive for CD31 (78,5%) and CD34 (82,2%) endothelial cell markers (Figure 8B). This was consistent with immunocytochemistry analysis showing von Willebrand factor (vWF) immunofluorescence in ECs cytoplasm and strong CD31 staining, visualizing the tight junctions between cells (Figure 8C). We coated microparticles with hiPSC-ECs and seeded them into arrays using our magnetic field-based micropatterning method. To ensure proper coating of the beads with hiPSC-ECs and to warrant cell adhesion to microbead surface, we pre-coated the microbeads with ACF Cell Attachment Substrate. During the first two days of culture, we observed intense sprouting of ECs from the beads followed by rapid invasion of individual ECs into the fibrin hydrogel (Figure 8D, 8E). The observations were qualitatively similar for two different sources of hiPSC-ECs. The hiPSC-ECs formed dense networks in the fibrin gel but failed to form mature tubular capillary structures.

**Figure 8.**
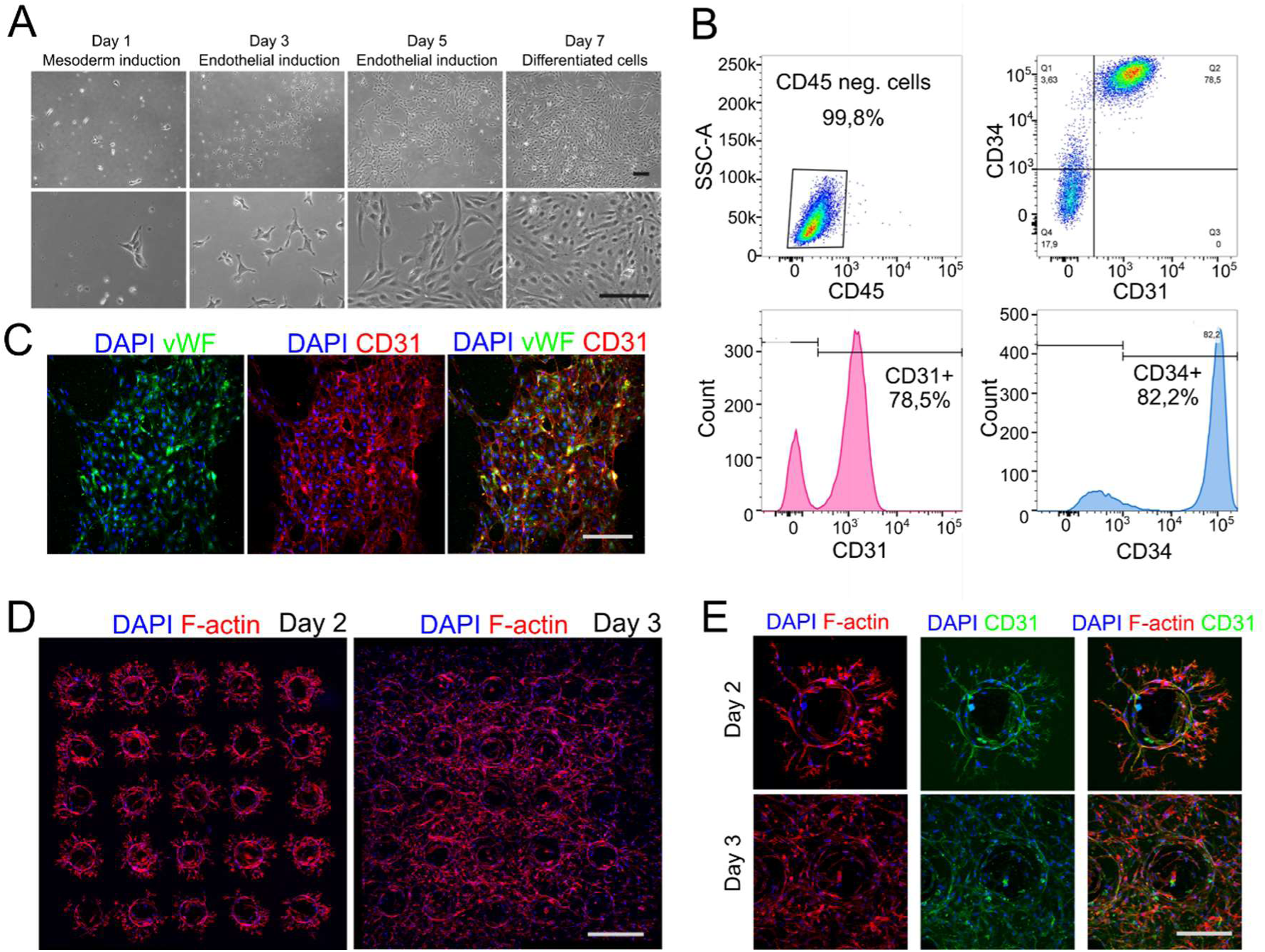
Generation of the arrays of hiPSC-ECs-coated microbeads. **A)** Bright-field images at different time points (day 1 to 7) of hiPSC-EC differentiation. Scale bar 25 µm. **B**) Flow-cytometry scatter plots and histograms of generated hiPSC-ECs stained for CD45, CD31 and CD34. **C**) Immunostaining of hiPSC-ECs for CD31 (red) and vVW factor (green). Hoechst dye (blue) was used to visualize nuclei. Scale bar 25 µm. **D**) Images of the arrays of hiPSC-EC-coated microbeads seeded at 500 µm spacing visualized at day 2 and 3 of culture. F-actin (red) was used to visualize the morphology of hiPSC-EC sprouts. Nuclei were stained with Hoechst dye (blue). Scale bar 500 µm. E) High magnification images of selected hiPSC-EC-coated microbeads immunostained for F-actin (red), CD31 (green) and Hoechst (blue) at day 2 and 3 of culture. Scale bar 250 µm.

We can therefore conclude that the behavior of the ECs, including their propensity towards the formation of monolayers or towards sprouting, may strongly depend on the specific cell type and origin (HUVECs vs hiPSC-ECs) [36]. Furthermore, hiPSC-ECs, unlike the well-established primary cells such as HUVECs, are strongly heterogeneous, which may result in different (e.g., patient-specific) angiogenic sprouting behavior [36]. Thus, to fully assess the potential of hiPSC-ECs in the generation of microvascular arrays and to possibly use such types of cells in drug tests warranting reproducible results, it is essential to optimize the endothelial cell differentiation protocols and source the representative cell batches from multiple donors. Such efforts are crucial for harnessing the full potential of hiPSC-ECs in developing reliable microvascular models for drug testing and personalized therapies.

## CONCLUSIONS

The magnetic field-driven pre-patterned microvascular arrays that we develop in this work open new perspectives in engineering of vascularized tissue models with broad applications ranging from regenerative medicine to drug discovery, including its potential use in development of personalized therapies. Our technology, based on the use of EC-coated superparamagnetic microbead-arrays, allows for fast formation of biologically mature and lumenized capillary networks of pre-designed architecture under minimal consumption of the endothelial cells.

Our approach might be of great value in terms of screening of the anti-angiogenic agents applied in cancer treatment or pro-angiogenic therapeutics that could support wound healing and tissue restoration therapies. The use of an ordered array of ‘angiogenic seeds’ warrants reproducible culture conditions (including spatial restrictions) whereas, at sufficiently large seed-seed spacing (in practice 1.5 mm or larger), each seed can be treated as an independent biological experiment. The effect of a drug can be monitored in terms of the resulting morphology of the vasculature quantified in terms of several metrics such as, e.g., (i) the area and length of the developed vascular network, (ii) the number of branches and their width (iii) the number of tips and (iv) the connectivity of microvasculature. The fully automated multiparametric morphometric analysis enables better understanding of the impact of the drugs on the tumor microvasculature, which would not be possible based on a single parameter (e.g., only the area of the sprouts) as typically employed in conventional angiogenesis assays [6, 44, 47, 48]. For example, our analysis does not show any significant impact of Taxol treatment on the area of the microvasculature, while the length and the number of vascular connections are significantly reduced and the width of the capillaries increases. Those type of effects were not reported in previous studies based on the sole microvascular area analysis in Taxol-treated samples [44]. Here we show that—only taken together— the multiple parameters give a full account of the changes in the morphometric phenotype of drug-treated capillaries. Furthermore, we find a significantly different morphological mechanism of action in the case of Sorafenib, i.e., predominantly capillary thinning and ‘overdose’ effects where increasing the drug concentration (from 1 µM to 10 µM in our case) leads to a quenched skeleton and unchanged or even elevated characteristics such as sprout length or the number of tips.

Moreover, having demonstrated the compatibility of the platform with hiPSC derived ECs, we strongly believe that it can be successfully applied in personalized drug screening studies in the future. Also, thanks to the open-top design of our culture chambers, the vascular microtissues can be easily extracted for use in further analysis including gene expression and/or spatial proteomics.

In summary, we have presented a novel vasculature-on-chip platform comprising human vascularized microtissues of highly organized and controlled architecture suitable for precise morphometric analysis of vascular networks and drug screening applications. We showed that the control over the initial architecture set by the distance between “vascular seeds” plays a critical role in the formation of microvascular networks and their final morphology. We demonstrated that the engineered vascularized microtissues grown in our platform are highly reproducible and the vasculature is functional (presence of lumens, tight junctions, etc.).

Our pre-patterned EC-microcarrier arrays allow for generation of mesoscale, spatially controlled microvasculatures of high biological relevance, and as such facilitate highly precise, reproducible morphological analyses including high-content phenotypic drug screens at the microtissue level.

## METHODS

### Device design and fabrication

The device was composed two layers. The bottom layer was 25 mm x 25 mm polycarbonate (PC) chip with embedded array of cylinder-shaped neodymium micromagnets grade N48, of diameter *D*_mag_ = 200 µm, *H*_mag_ = 500 µm. The upper layer was a 30 mm × 30 mm PDMS chip (SYLGARD™ 184 Silicone Elastomer Kit, Dow Corning Corporation, Michigan, United States) with a culture chamber of the dimensions 10 mm × 10 mm bonded to a 24 mm × 24 mm #0 glass coverslip (85 - 115 µm µm; ISO 8255-1:2017) (THORLABS, Mölndal, Sweden; catalog no. CG00C2) as the bottom wall. Before bonding, the glass coverslip and the PDMS chip were rinsed in 70% ethanol and dried under the biological safety cabinet. To accommodate an array of neodymium cylinder-shaped micromagnets in the PC chip, first, the array of holes was fabricated on its surface using CNC micromilling according to the chosen design. Subsequently, the PC chip was placed on top of a large neodymium magnet. The chip was observed under a stereoscopic microscope and micromagnets were manually positioned inside the holes using non-magnetic titanium fine tip tweezers. The spacing between the micromilled holes determined the spacing between the neodymium micromagnets. During the magnetic assembly of EC-microcarriers the device was assembled in such a way that the PDMS chip was mounted on top of the PC chip, tightly aligned with UHU patafix paste to avoid any displacements, and placed under the stereoscopic microscope for inspection and possible corrections.

### Numerical Simulations

Numerical simulations were performed using COMSOL Multiphysics 6.1 with the AC/DC Module. The simulations were based on the “Magnetic Fields, No Currents” interface to model the magnetic field distributions. The program solves Maxwell’s Equation ∇⋅**B** = 0, where **B** is the magnetic field defined as **B** = μ_0_μ_r_**H** + **B**_r_, where μ_0_ and μ_r_ correspond to vacuum and relative permeabilities, **B**_r_ is remanent magnetic field, and **H** is the magnetic field strength. Values of μ_r_ for materials used in this study, i.e., air, NeFeB, and iron, are 1, 1.05, and 4000, respectively. For the domains not being sources of the field, **B**_r_ is 0. For the domains acting as permanent magnets, **B**_r_ was set to 1.4 T, a value corresponding to N48 grade magnets. **H** is defined as **H** = -∇V_m_, where V_m_ is the magnetic scalar potential. A zero-flux boundary condition **n**⋅**B** = 0 was set on the external edges of the system (see Figure S1). Two geometries were constructed for the study, as depicted in Supplementary Figure 6.

Geometry 1: This model simulated the magnetic field distribution over a ferromagnetic wire with a diameter of 200 μm and a height of 500 μm. A cylindrical permanent magnet (20 mm diameter × 5 mm height) was placed beneath the wire. The permanent magnet served as the source of the magnetic field, while the wire acted as a conduit to guide and concentrate the magnetic field over a localized area.

Geometry 2: This model simulated a permanent NeFeB micromagnet (with dimensions of the wire used in the previous configuration) that generates a magnetic field autonomously.

Both geometries were constructed using a 2D axisymmetric representation to optimize computational efficiency and accurately model the physical setup. To improve the accuracy of the solution in the vicinity of the magnetized wire or micromagnet we introduced an additional fine meshed air domain.

### Cell culture

GFP-HUVECs and RFP-HUVECs (Angio-Proteomie, Boston, MA, USA; catalog no. cAP-0001GFP and cAP-0001RFP) were cultured in Endothelial cell growth medium 2 (EGM-2) medium supplemented with EGM-2 bulletkit (Lonza, Basel, Switzerland; catalog no. CC-3156 & CC-4176) and were used at passages 3 through 5. NHDF (Promocell, Heidelberg, Germany; catalog no. C-12302) were cultured in Dulbecco’s modified Eagle medium (DMEM) (Thermo Fisher Scientific, Waltham, MA, USA; catalog no. 10566016) supplemented with 10% fetal bovine serum (FBS), Glutamax, 1% penicillin-streptomycin. NHDF were used between passages 2 and 7. GFP-HeLa cells (GeneTarget Inc, San Diego, CA, USA; catalog no. SC034-Puro) were cultured in DMEM supplemented with 10% FBS, Glutamax, 1% penicillin-streptomycin and 0.1 mM Mem Non-Essential Amino Acid Solution (Merck KGaA, Darmstadt, Germany catalog no.7145). All cells were cultured in 5% CO_2_ at 37 °C humidified atmosphere and media were replaced every 2 days.

### hiPSC culture and differentiation into endothelial cells

Commercial hiPSC lines derived from healthy donors (ThermoFisher Scientific, Waltham, MA, USA; catalog no. A18944) were cultured and expanded using TeSR E8 complete medium (StemCell Technologies, Vancouver, BC, Canada; catalog no. 05990 and 05991) at 37°C, 5% CO_2_ and closely monitored daily. The Endothelial Differentiation Kit (StemCell Technologies, Vancouver, BC, Canada; catalog no. 08005) was employed for the iPSC differentiation into endothelial cells (EC). Briefly, iPSCs were dissociated with Gentle Cell Dissociation Reagent (StemCell Technologies, Vancouver, BC, Canada; catalog no. 07174) at 37°C for 8 min. A single-cell suspension was obtained pipetting up and down 3 - 4 times and the cells were collected and centrifuged at 300*g* for 5 min. The hiPSCs were then resuspended in fresh Tesr-E8 medium supplemented with 10 µM RevitaCell (Thermo Fisher Scientific, Waltham, MA, USA; catalog no. A2644501) and 2×10^5^ cells/well were seeded on pre-coated Matrigel (Corning, NY, USA; catalog no. 354277) 6-well plates. After 24 hours of incubation at 37°C, the medium was replaced with STEMdiff Mesoderm Induction Medium (StemCell Technologies, Vancouver, BC, Canada; catalog no. 05221). Starting from day 3, cells were maintained in STEMdiff Endothelial Induction Medium (StemCell Technologies, Vancouver, BC, Canada; catalog no. 08006), and the medium was changed every 48 h. By day 7, cells were enzymatically detached using Accutase (StemCell Technologies, Vancouver, BC, Canada; catalog no. 07920) and seeded on 6-well plates pre-coated with ACF Cell Attachment Substrate (StemCell Technologies, Vancouver, BC, Canada; catalog no. 07130) in complete STEMdiff Endothelial Expansion Medium (StemCell Technologies, Vancouver, BC, Canada; catalog no. 08008 and 08009). A seeding density of 1×10^4^ cells/cm^2^ was used to achieve cell confluency between days 3 and 5 post-seeding. Immunofluorescence and flow cytometry were performed on day 7 of the protocol to evaluate the EC differentiation.

### Coating of the beads with endothelial cells

Coating of the beads with HUVECs was performed as described previously [12]. Briefly, fluorescently tagged HUVECs were mixed with monodispered polystyrene superparamagnetic microcarrier beads of diameter 265µm, SD = 3.4µm (CV = 1.3 %; microParticles GmbH, Berlin, Germany; catalog no. PS-MAG-AR111) at the concentration of approximately ∼ 500 cells per bead in a small volume of warm EGM-2 medium and placed in the incubator for 4 hours at 37°C and 5% CO_2_, gently shaking the tube every 20 min. After 4 hours, beads were transferred to a culture flask with fresh EGM-2 medium and placed overnight in the incubator at 37° C and 5% CO_2_. The following day beads were seeded in a culture chamber of the device.

For hiPSC-EC microcarrier beads were pre-coated with ACF Cell Attachment Substrate (StemCell Technologies, Vancouver, BC, Canada; catalog no. 07130) for 2h at RT with agitation. Subsequently, the beads were transferred to a new Eppendorf tube, washed 3 times with PBS for 5 min, resuspended in complete STEMdiff Endothelial Expansion Medium (StemCell Technologies, Vancouver, BC, Canada; catalog no. 08008 and 08009) and mixed with hiPSC-EC for 2 hours at 37° C and 5% CO_2_, gently shaking the tube every 20 min. After 2 hours, beads were transferred to a culture flask with fresh complete STEMdiff Endothelial Expansion Medium (StemCell Technologies, Vancouver, BC, Canada; catalog no. 08008 and 08009) and placed for 2 hours in the incubator at 37° C and 5% CO_2_. The same day hiPSC-EC coated beads were seeded in the culture chamber of the device.

### Seeding the device and generation of the arrays

EC-coated beads were gently washed with fresh medium, appropriate for the used cells, and resuspended in freshly prepared 2.5 mg/ml fibrinogen solution (Sigma-Aldrich, St. Louis, MO, USA; catalog no. 341573). Precisely, for each array, 25 microbeads were resuspended in 180 µl of 2.5 mg/mL fibrinogen solution, and poured into the culture chamber of the PDMS chip. For HUVECs the fibrinogen solution was additionally mixed with 100 000 hDFs. The role of hDFs was to enhance the angiogenic behavior of HUVECs. After positioning the PDMS chip containing the polystyrene microparticles above the PC chip containing the array of cylinder-shape neodymium micromagnets, fibrinogen solution was aspirated to an Eppendorf tube, mixed with 20 µl of 6.25U/mL thrombin solution (Sigma-Aldrich, St. Louis, MO, USA; catalog no. T4648) and poured back into the culture chamber of the PDMS chip allowing hydrogel the hydrogel to crosslink (1-2 minutes). The array generation process consisted of two stages. Immediately after pouring the hydrogel solution containing the microparticles into the PDMS culture chamber, approximately the 80% of microparticles self-assembled over the magnetic hotspots. The remaining microparticles were manually positioned using non-magnetic titanium fine-tip tweezers. Fibrin/bead solution was allowed to clot for 5 minutes at room temperature and then at 37° C and 5% CO_2_ for 30 minutes. Finally, 0.8 mL of medium was added to the culture chamber of each PDMS chip. For HUVECs complete EGM-2 supplemented with 10 ng/mL human recombinant VEGF-165 (StemCell Technologies, Vancouver, BC, Canada; catalog no.78073) was used. Arrays of hiPSC-EC-coated beads were cultured in complete STEMdiff Endothelial Expansion Medium (StemCell Technologies, Vancouver, BC, Canada; catalog no. 08008 and 08009) without heparin. The medium was changed every other day.

### Drug exposure studies

Arrays of RFP-tagged HUVECs were cocultured with NHDFs and GFP-tagged Hela cells to mimic the cancer microenvironment. The ratio of NHDFs and HeLa cells was 1:10. For screening assays, after 4-5 days of culture arrays were exposed to 100 nM and 1000 nM Taxol (Merck KGaA, Darmstadt, Germany, catalog no. PHL89806) or 1 µM and 10 µM Sorafenib (Merck KGaA, Darmstadt, Germany, catalog no. SML2653) for 48 h. The morphometric analyses were performed on days 4-5 before adding a drug, and after 48 h of exposure. Compounds were dissolved in dimethyl sulfoxide (DMSO) and added to the medium with less than 0.01% DMSO. The controls were treated with the appropriate amount of DMSO.

### FACS analysis

hiPSC-derived ECs were detached, washed, and incubated at RT for 20 min with the following fluorochrome-conjugated antibodies: CD34 Pe-Cy5 (Beckman Coulter, Brea, CA, USA; catalog no. A07777), CD31 BUV737 (BD Biosciences, Franklin Lakes, NJ, USA; catalog no. 748320), and CD45 BV421 (Biolegend, San Diego, CA, USA; catalog no. 368521). Then, cells were washed and resuspended in PBS supplemented with 5% FBS and 2 mM EDTA. The samples were acquired by BD FACSymphony A5 instrument and data were analyzed by FlowJo software v10.8.1.

### Immunofluorescence staining

Cells and fibrin blocks were fixed with 4% paraformaldehyde (PFA) for 15 min. Samples were blocked for 30 min with blocking buffer (2% bovine serum albumin [BSA], 2% normal goat serum, and 0.5% Triton X-100 in PBS) at room temperature before overnight incubation at 4°C with primary antibodies. The following primary antibodies were used: anti-CD31 (Abcam, Cambridge, MA, USA; catalog no. ab24590), anti-Von Willebrand factor (Abcam, Cambridge, MA, USA; catalog no. ab11713), anti-laminin (Sigma-Aldrich, St. Louis, MO, USA; catalog no. L9393) . The following day, the samples were washed 3 times for 5 min in PBS and incubated with appropriate fluorescent-conjugated secondary antibodies for 2 h at RT. Primary and secondary antibodies were applied in the blocking buffer. Alexa Fluor 568 Phalloidin (Thermo Fisher Scientific, Waltham, MA, USA; catalog no. A12380) was applied together with secondary antibodies. To visualize nuclei Hoechst dye 2 mg/ml (Thermo Fisher Scientific, Waltham, MA, USA; catalog no. H21491) was used at working dilution 1:1000.

### Isolation of microvascular arrays from the chip

To isolate microvascular arrays, the fibrin blocks containing microparticles were first washed with PBS and then PDMS chip was disassembled manually in such a way that the fibrin block with the array of microparticles remained on the surface of the coverslip. Arrays were transferred into PBS-containing Petri dish using a laboratory spatula.

### Image acquisition and analysis

Fluorescence microscopy was performed using Nikon A1 confocal microscope (Nikon Instruments, Inc, Melville, NY, USA) equipped with a PLAN APO 10×/0.45 or 60×/1.20 objectives. Images were collected using NIS-Elements Advanced Research software (Nikon Instruments, Inc, Melville, NY, USA) in the nd2 format, in 16-bit single-channel, with a typical resolution of 1.25 µm/pix with respective metadata.

For the morphometric analysis of microvascular networks, we modified a previously developed Python-based segmentation protocol dedicated to morphometric analysis of a single sprouting microbead [12] to operate in a system composed of multiple microbeads. The segmentation pipeline followed the workflow illustrated in Figure 2B. Briefly, the images were first max-pooled on the Z-directional slices (taking the maximum intensity value across the stack) and treated with a Gaussian blur followed by adaptive thresholding using the OpenCV2 library [49]. After the generation of a binary mask representing endothelial cells, two morphological operations—closing and opening—were performed to connect elements from the adaptive threshold output and to eliminate noise. Next, an additional smoothing step was performed, filling small holes of perimeter smaller than 250 µm. Finally, some disconnected elements of the network were removed which amounted to 3 % of the total area, thus almost eliminating any background noise while not significantly affecting the network morphology.

Next, the centers of microcarriers were identified. Part of this task was performed manually, and the results were used to train and apply the YOLO v7 machine learning architecture [50]. The trained software automatically detected the remaining microbead centers in our datasets. The final outputs were reviewed and corrected manually, if necessary. The binarized images were then skeletonized and masked with circles corresponding to the locations of the micro-carriers. Processed images were then used to generate a graph representation, allowing for the calculation of morphological metrics including the total area of the sprouts (i.e., with the masked areas excluded), the total length of the sprouts, the number of tips (connected to at least one micro-carrier), the number of connected components, and the average width of the sprouts (equal to the total area divided by the total length).

Connected component (CC) can be defined as a group of connected bright pixels, with each pixel having in its Moore neighborhood another bright pixel, where Moore neighborhood is defined as the 8 pixels adjacent to a given pixel. In the case of microbead arrays, we limited the definition of CC only to the groups of connected pixels whose area was comparable to the area of a single sprouting bead, *A*_1_ = *A*_tot_/*N*, where *A*_tot_ is the total area of the network (including the area of the cells at the beads) and *N* = 5 × 5 = 25. We only counted those CCs that fulfill the condition for their area: *A*_CC_ > 0.6 *A*_1_.

Additionally, we evaluated network connectivity by counting the number of intersections between the network (excluding the tip segments) and the Voronoi cells centered at the microcarriers (Figure 2C). Voronoi division was also employed to assess the directionality of growth of the capillaries around the individual microcarriers. For each Voronoi cell, and thus for each microcarrier, we measured mass distribution in octants aligned with the directions of a wind rose. The micro-carrier population was classified based on the number of nearest neighbors: corner (3 neighbors), bar (5 neighbors), and center (8 neighbors). Asymmetrical results for corner and bar regions were transformed to a common coordinate system, with all regions oriented to the right (e.g. top bar is transformed by -90 degrees, etc.).

### Analysis of the angular area distribution

The circular histograms (radar-charts) presented in Figure 3A and Supplementary Figure 2 and 3 consist each of eight bins that evenly divide the range (−π,π) in the polar coordinate system. The radius of each circular ‘slice’ was calculated so that its area was proportional to the vascular network mass in the corresponding angular range. The areas for a given day were normalized by the total area of the vascular network on day 7 averaged over the group of beads with the corresponding location within the matrix (center, edges, or corner). This allowed us to capture the evolution of angular mass distribution with respect to some reference point.

Coefficient of variation of the angular area distribution (Figure 3B) was calculated as the standard deviation of the areas of the octants for each day, divided by its mean value.

At small bead-to-bead distances (*d* = 400, 500 µm), where the network becomes dense, the histograms calculated using the above method may reflect the shape of Voronoi cells rather than the rate of sprouting. This is because the shape of the Voronoi cells is close to a square so that the octagonal division results in Voronoi-cell subsections of different areas (see Figure 2C). Therefore, to address the rate of sprouting more precisely, we also calculated a normalized histogram (for the case *d* = 500 µm at day 7), in which the network area within each octant was divided by the area of the corresponding Voronoi-cell subsection. We applied this approach only for the central beads (3 x 3 subarray), where the Voronoi cells were indeed nearly square-shaped (see Supp. Figure 3).

### Statistical analysis

Directly measured quantitative data were expressed as the mean ± SEM. The statistical methods (one-way analysis of variance followed by Tukey’s post hoc test, and two-way analysis of variance followed by Bonferoni post hoc test, two-tailed unpaired t-test) and *p*-values were defined in the figure legends or supplementary material Table 1. The statistical analyses were performed using GraphPad Prism 5 and Microsoft Excel software. The results of the statistical analysis are available in the following sheets of the Supp. Table1. Numerical values that underlie the graphs are shown in Supp. Data file.

## SUPPORTING INFORMATION

For Supporting Information, please see Supplementary Figures file and Supplementary video 1 and 2. The results of the statistical analysis are available in Supplementary Table1. Numerical values that underlie the graphs are shown in Supplementary Data file.

## ACKNOWLEDGEMENTS

Preparation of this article was supported by Sonatina (Grant No. 2020/36/C/NZ1/00238 awarded to K.O.R.) and Opus (Grant No. 2022/45/B/ST8/03675 awarded to J.G.) from the Polish National Science Center (NCN).

## DATA AVAILABILITY STATEMENT

The data that support the findings of this study are available in the supplementary material of this article.

## CODE AVAILABILITY STATEMENT

Code is available to upon request.

