## Supplementary Figures for "Engineering of interconnected microcapillary networks at the mesoscale via magnetic assembly of endothelial-cell ‘seeds’"

### Magnetically assembled arrays of endothelial-cell microcarriers as a versatile tool in vascular tissue engineering, drug testing and angiogenesis research.

**Katarzyna O. Rojek [1], Antoni Wrzos [2], Fabio Maiullari [1,3], Konrad Giżyński [1], Maria Grazia Ceraolo [4], Claudia Bearzi [3], Roberto Rizzi [5], Piotr Szymczak [2] and Jan Guzowski [1*]**

1. Institute of Physical Chemistry, Polish Academy of Sciences, Kasprzaka 44/52 , 01-224 Warsaw, Poland
2. Institute of Theoretical Physics, Faculty of Physics, University of Warsaw, Pasteura 5, 02-093 Warsaw, Poland
3. Institute for Biomedical Technologies, National Research Council, Via Fratelli Cervi, 93, Segrate, 20054 Milan, Italy
4. Neurology Unit, Fondazione IRCCS Ca' Granda Ospedale Maggiore Policlinico, 20122 Milan, Italy
5. Department of Medical-Surgical Sciences and Biotechnologies, Sapienza University, C.so della Repubblica 79, 04100 Latina, Italy

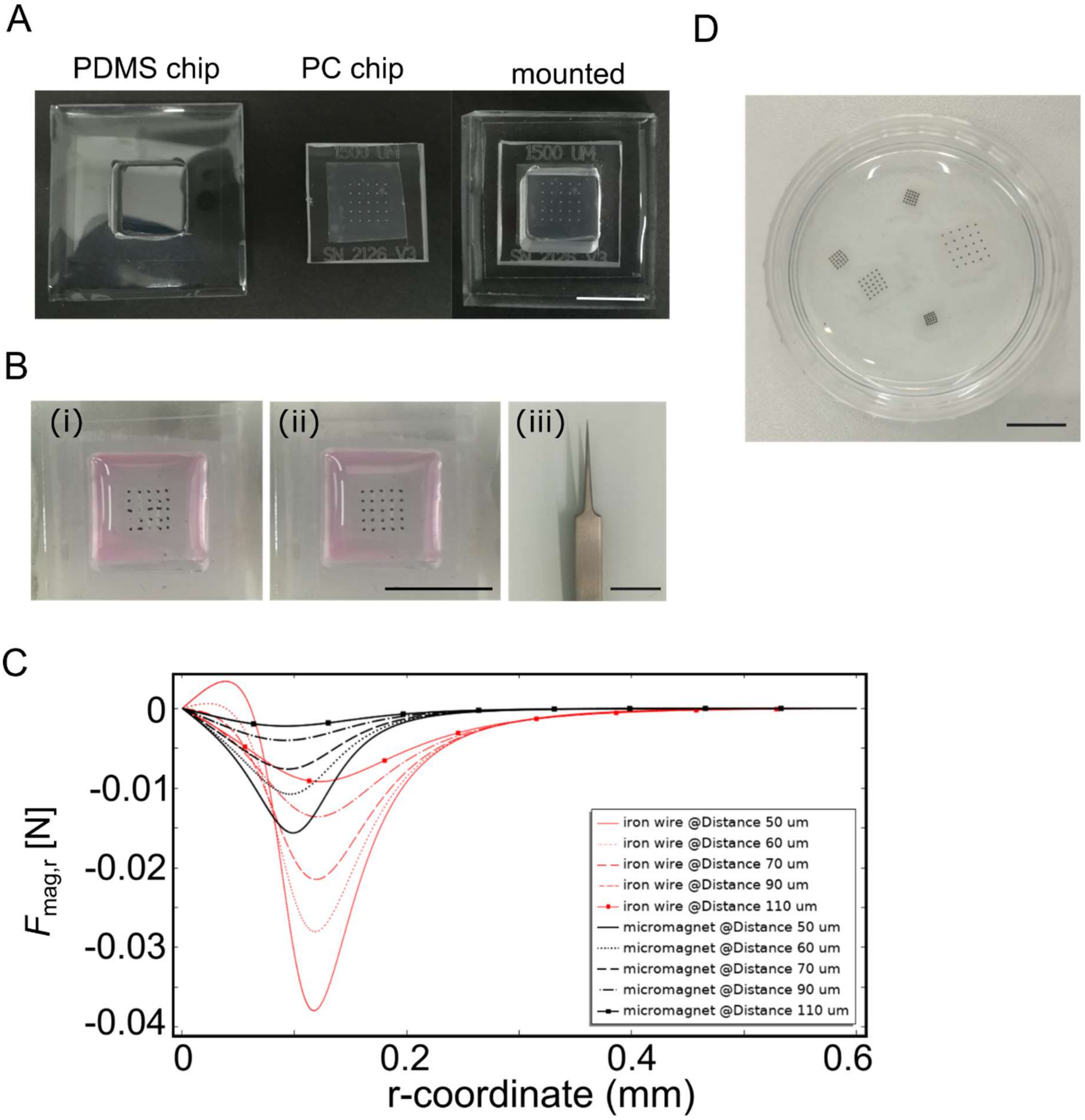


**Supplementary Figure 1.** Generation of ordered arrays of microparticles over the substrate using magnetic field patterning. **A**) Images of PDMS culture chamber, PC chip with embedded micromagnets and the assembled device. **B**) The magnetic microbeads that did not spontaneously migrate to the magnetic-field hotspots can be manually corrected using titanium fine-tip tweezers. **C**) Results of the numerical calculation of the magnetic force $F_{\mathrm{mag,r}}$ acting at a probe microparticle depending on its radial displacement $r$ from the micromagnet axis for the experimentally relevant spacing $H = 90$ µm. Positive direction of the force is outward (i.e., negative values correspond to magnetic attraction). **D)** Isolated hydrogel sheets with embedded arrays of EC-coated microparticles in PBS in P35 Petri dish. Scale bars 1 cm.

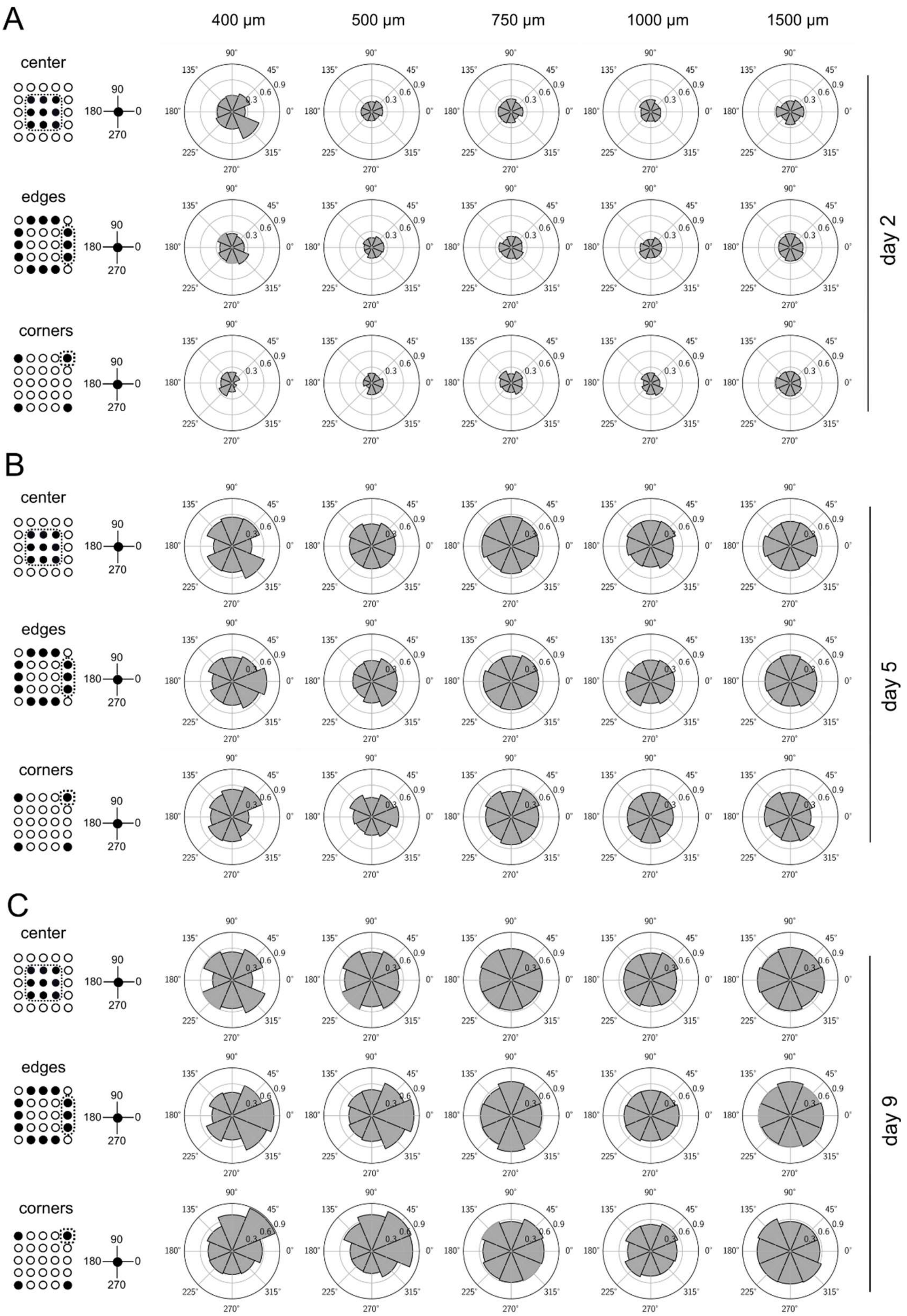


**Supplementary Figure 2.** Directionality analysis of the sprouting microcapillary networks. **A-C**) Radar charts illustrating the angular (azimuthal) distribution of the area occupied by capillaries at day (**A**) 2, (**B**) 5 and (C) **9** of culture. EC-coated microbeads were classified into 3 groups based on the number of nearest neighbors: center (8 neighbors), edge (5 neighbors) and corner (3 neighbors). Asymmetrical results from corner and edge regions were transformed to a common coordinate system, with all regions oriented to match the region indicated with a dotted frame (eg. top bar is transformed by -90 degrees, etc.). The numbers of the arrays in the assay (biological repetitions) were as follows: Spacing 400 µm, *n* = 3; 500 µm, *n* = 5; 750 µm, *n* = 7; 1000 µm, *n* = 6; 1500 µm, *n* = 7).

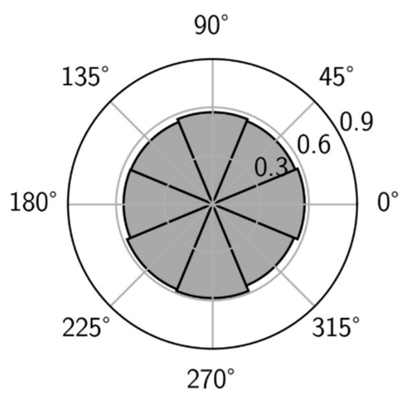


**Supplementary Figure 3.** Directionality analysis of the sprouting microcapillary networks formed around the central beads (3 x 3 subarray) seeded at $d$ = 500 µm. Radar chart illustrating the angular (azimuthal) distribution of the area occupied by capillaries at day 7 of culture normalized by the area of the Voronoi-cell subsection in each octant. The number of the arrays in the assay (biological repetitions) $n$ = 5.

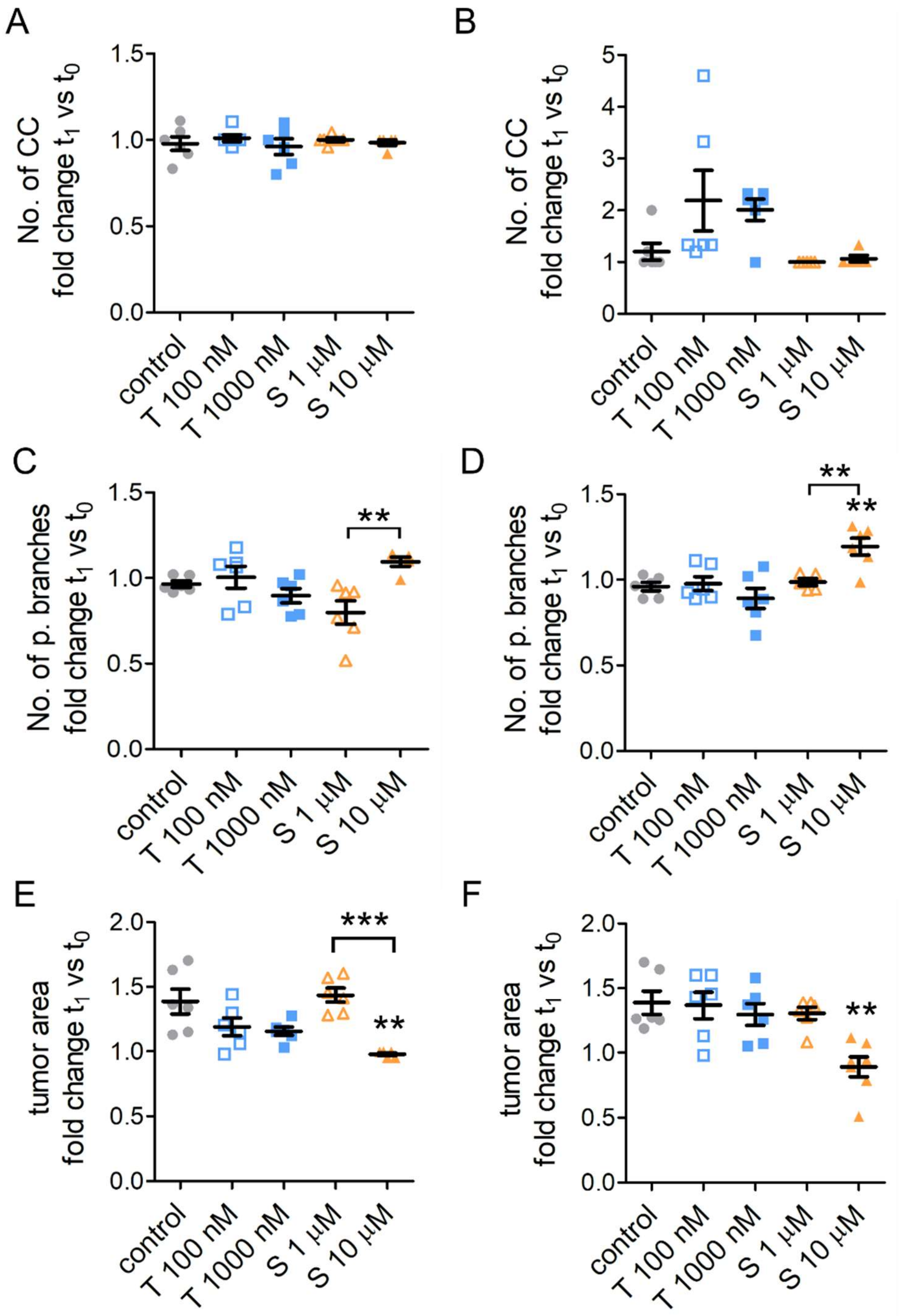


**Supplementary Figure 4.** Morphometric analysis of the fold-change in (**A,B**) the number of connected components (CC), (**C, D**) the number of primary branches and (**E,F**) the tumor area at time $t_1$ relative to $t_0$ for the arrays of HUVEC-coated microbeads with **(A, C, E)** $d$ = 1500 µm and **(B, D, F)** $d$ = 500 µm spacing co-cultured with HeLa cells and exposed to anticancer drugs. Spacing 500 µm: Control, $n$ = 6; Taxol (T) 100 nM, $n$ = 6; Taxol (T) 1000 nM, $n$ = 6; Sorafenib (S) 1 µM, $n$ = 5; Sorafenib (S) 10 µM, $n$ = 6. Spacing 500 µm: Control, $n$ = 6; Taxol (T) 100 nM, $n$ = 6; Taxol (T) 1000 nM, $n$ = 6; Sorafenib (S) 1 µM, $n$ = 6; Sorafenib (S) 10 µM, $n$ = 5. Error bars around the mean indicate the standard error of the mean (SEM). * $P < 0.05$, ** $P < 0.01$, *** $P < 0.001$. Statistical significance was analyzed using one-way ANOVA with Turkey post hoc test. Numerical values that underlie the graphs are shown in Supp. Data file.

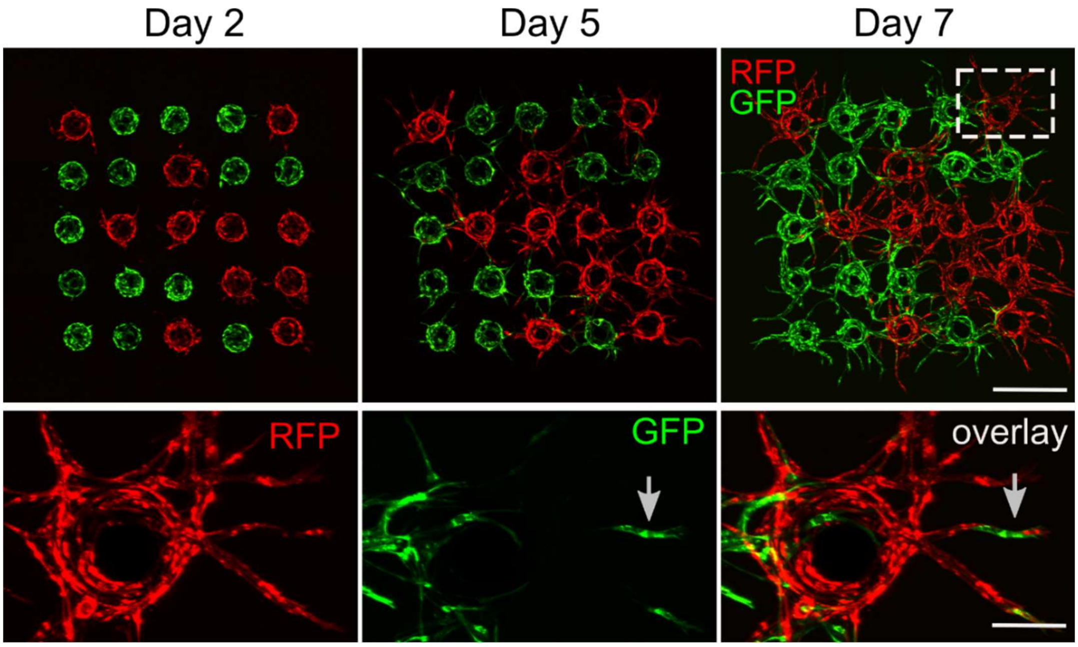


**Supplementary Figure 5.** An array of GFP and RFP-tagged HUVEC-coated microbeads seeded at $d$ = 500 µm and imaged at days 2, 5 and 7 of culture. Bottom panel shows high magnification images of the selected area. Arrow indicates a single GFP tagged cell that migrated from the GFP-HUVEC coated microbead to the distal part of RFP-HUVEC coated microbead. Scale bar 500 µm (upper panel) and 100 µm bottom panel.

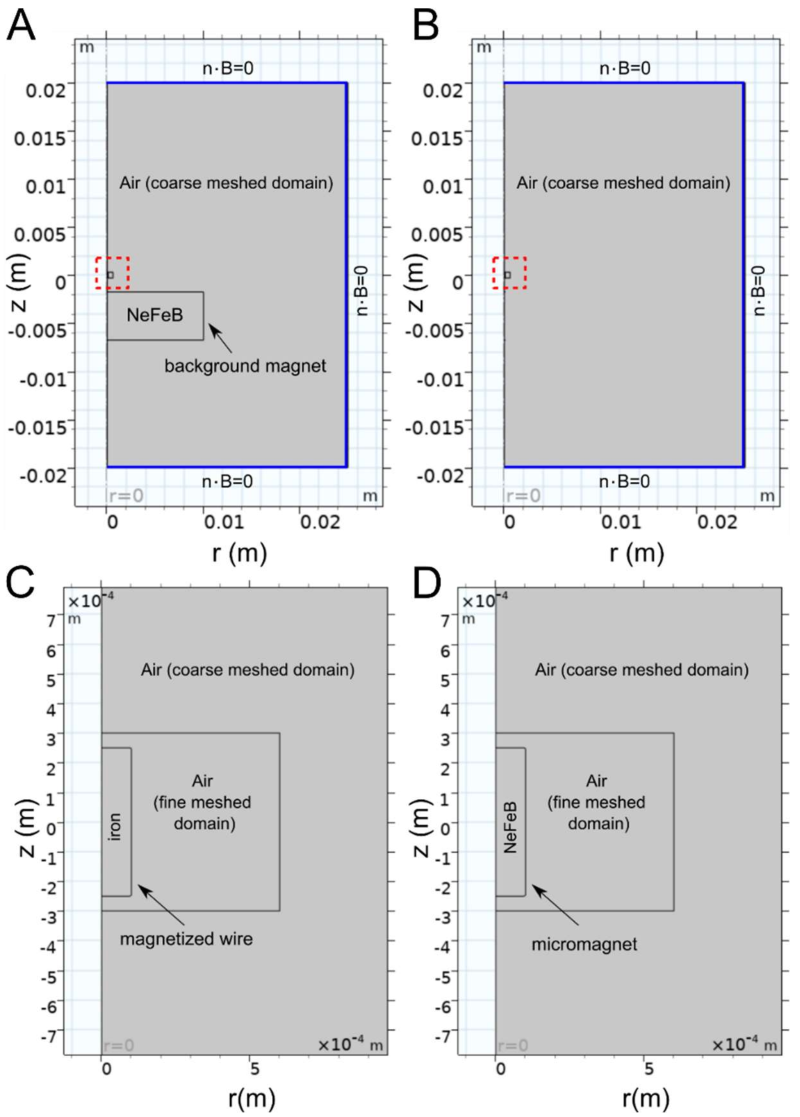


**Supplementary Figure 6**. Schematic representation of (**A**) Geometry 1 and (**B**) Geometry 2 showing the air (coarse meshed domain), and the region of interest (red dashed box). At the external boundaries of the system (marked with blue color), a 50 µm geometry layer was added to serve as an artificial domain with a physical width equal to 47 m. In (**A**) there is a background magnet placed beneath ROI. (**C**,**D**) Zoom-in of ROI (**A**) and (**B**) respectively. Air domain around the micromagnet/iron wire is meshed with smaller elements.
